# NEBULA: a fast negative binomial mixed model for differential expression and co-expression analyses of large-scale multi-subject single-cell data

**DOI:** 10.1101/2020.09.24.311662

**Authors:** Liang He, Alexander M. Kulminski

**Author notes:** Corresponding authors: Liang He, Alexander Kulminski.

## Abstract

The growing availability of large-scale single-cell data revolutionizes our understanding of biological mechanisms at a finer resolution. In differential expression and co-expression analyses of multi-subject single-cell data, it is important to take into account both subject-level and cell-level overdispersions through negative binomial mixed models (NBMMs). However, the application of NBMMs to large-scale single-cell data is computationally demanding. In this work, we propose an efficient NEgative Binomial mixed model Using a Large-sample Approximation (NEBULA)), which analytically solves the high-dimensional integral in the marginal likelihood instead of using the Laplace approximation. Our benchmarks show that NEBULA dramatically reduces the running time by orders of magnitude compared to existing tools. We showed that NEBULA controlled false positives in identifying marker genes, while a simple negative binomial model produced spurious associations. Leveraging NEBULA, we decomposed between-subject and within-subject overdispersions of an snRNA-seq data set in the frontal cortex comprising ∼80,000 cells from a cohort of 48 individuals for Alzheimer’s diseases (AD). We observed that subpopulations and known subject-level covariates contributed substantially to the overdispersions. We carried out cell-type-specific transcriptome-wide within-subject co-expression analysis of *APOE*. The results revealed that *APOE* was most co-expressed with multiple AD-related genes, including *CLU* and *CST3* in astrocytes, *TREM2* and C1q genes in microglia, and *ITM2B*, an inhibitor of the amyloid-beta peptide aggregation, in both cell types. We found that the co-expression patterns were different in *APOE2*^+^ and *APOE4*^+^ cells in microglia, which suggest an isoform-dependent regulatory role in the immune system through the complement system in microglia. NEBULA opens up a new avenue for the broad application of NBMMs in the analysis of large-scale multi-subject single-cell data.

## Introduction

Single-cell profiling technology has revolutionized our understanding of biology to an unprecedented resolution. With the recent advances in high-throughput single-cell profiling techniques (Hashimshony et al., 2016; Klein et al., 2015, 2015; Picelli et al., 2014), large-scale multi-subject single-cell RNA-seq (scRNA-seq) and ATAC-seq (scATAC-seq) data become increasingly accessible. Notably, through droplet-based technology, tens of thousands of cells from multiple subjects can be sequenced in parallel within a single batch (Klein et al., 2015; Macosko et al., 2015). As an example, a recent large-scale population single-nucleus RNA-seq (snRNA-seq) data in the human frontal cortex involving ∼80,000 nuclei from a cohort comprising 24 Alzheimer’s disease (AD) patients and 24 healthy controls (Mathys et al., 2019) provide biological information in much finer granularity compared to bulk RNA-seq data. Projects profiling more than one million single cells from hundreds of subjects are underway.

The drastically increasing magnitude of sample size, however, poses a significant computational challenge for efficient analysis of differential expression, expression quantitive trait loci (eQTLs), and co-expression using large-scale scRNA-seq data. This is in contrast to previous statistical models for analyzing bulk RNA-seq data, in which more emphasis is placed upon building a robust estimate under a small sample size (e.g., regularization of standard errors and robust estimation of overdispersion parameters (Law et al., 2014; Love et al., 2014; McCarthy et al., 2012)). Unlike bulk RNA-seq data, multi-subject scRNA-seq data, particularly using the droplet-based technology, often comprise many cells but are featured by high sparsity and a hierarchical structure. Due to the hierarchical nature of multi-subject scRNA-seq data, it is preferable to decompose the overdispersion into the subject and cellular levels. One of the standard approaches to handle hierarchical count responses is a negative binomial mixed-effects model (NBMM) that introduces independent random effects to take into account the overdispersion at both levels. In this paper, we focus on the NBMM rather than a zero-inflated model because multiple recent studies show that a zero-inflated model might be redundant for unique molecular identifiers (UMI)-based single-cell data (Chen et al., 2018; Choi et al., 2020; Hafemeister and Satija, 2019). This is also consistent with the observations for the vast majority of genes in our analysis of real data. The NBMM can increase the statistical power by a more accurate specification of the overdispersion structure, and also helps in eliminating spurious associations to detect marker genes of a cell cluster, as we show in our simulation.

A plethora of estimation methods have been proposed for such a model (Breslow and Clayton, 1993; Lindstrom and Bates, 1990; Ormerod and Wand, 2012; Rue et al., 2009; Tierney and Kadane, 1986; Zhang et al., 2017), and it is the invention of these fast algorithms that popularizes its practical use. However, the computational efficiency of existing tools (Bates et al., 2014; Brooks et al., 2017; Rue et al., 2009) still cannot suffice for the broad application in large-scale multi-subject scRNA-seq data. Due to the intractable marginal likelihood in the NBMM, current estimation algorithms generally rely on a two-layer iteration procedure, which often converges slowly. To address the computational burden, we propose a **NE**gative **B**inomial mixed model **U**sing **L**arge-sample **A**pproximation (NEBULA), a novel fast algorithm for association analysis of scRNA-seq data using an NBMM. As the core idea in NEBULA, we developed an analytical approximation of the high-dimensional integral for the marginal likelihood of the NBMM, by leveraging a large number of cells per subject (CPS), which is a common feature in droplet-based scRNA-seq data. The improvement of computational efficiency is achieved by avoiding high-dimensional optimization and converging within fewer steps.

We demonstrated the efficiency and accuracy of NEBULA through an extensive simulation study. Because NEBULA uses approximation based on large numbers, we explored the questions of how many cells and subjects are required to estimate subject-level and cell-level overdispersions accurately. We also investigated the performance of NEBULA in dealing with low-expressed genes, and controlling the false positives for testing fixed-effects predictors. It should be noted that a predictor of interest can be classified as a subject-level or a cell-level variable. The first category often includes disease or treatment groups, genetic variants, and subject-level covariates such as age, sex, etc, while stages of the cell cycles, subpopulations, and cell-level expression are common variables in the second category. For testing a cell-level variable, it is known that ignoring the subject-level random effects would lead to inflated p-values (Milanzi et al., 2012). In contrast, there is much less literature on the performance of testing subject-level predictors in cell-level data. It is well known that the subject-level random effects need to be included in the model when the predictor of interest is also a subject-level variable. Otherwise, the type I error rate would be inflated due to an underestimated standard error of the effect size. A detailed derivation of such underestimation is given in (Landeghem et al., 2005; Moerbeek, 2004). In this study, we found that testing a subject-level variable was highly sensitive to the estimation of the subject-level variance component. Thus, testing predictors at the two levels might require different estimation strategies.

To date, few studies have investigated the relative contributions of subject-level and cell-level overdispersions in cell-type-specific gene expression. Overdispersion often results from failing to include important explanatory variables or a hierarchical structure in the data (Hilbe, 2011). Using NEBULA, we decomposed gene-specific overdispersion of a large-scale snRNA-seq data in the human frontal cortex from an AD cohort (Mathys et al., 2019). We further explored what factors might contribute to the overdispersions. We showed that NEBULA reduced false positives in selecting cell-type marker genes, especially when the numbers of cells across subjects were unbalanced. As a direct application of NEBULA, we performed a cell-level transcriptomic co-expression analysis of *APOE* to investigate its cell-type-specific regulatory mechanisms in microglia and astrocytes. We further carried out an isoform-specific co-expression analysis for different *APOE* genotypes separately.

## Results

### Overview of NEBULA

The variation of raw counts in scRNA-seq data comes from two sources, the Poisson sampling process and overdispersion originating from biological heterogeneity and technical or experimental noise. Cells in large-scale single-cell data are often collected from multiple subjects or batches, leading to an apparent hierarchical structure. To model this strucutre, NEBULA decomposes the total overdispersion into subject-level (i.e., between-subject) and cell-level (i.e., within-subject) components using a random-effects term parametrized by *σ*^2^ and the overdispersion parameter *ϕ* in the negative binomial distribution. The subject-level overdispersion captures the contributions from e.g., eQTLs, technical artifacts, batch effects, and other subject-level covariates. The cell-level overdispersion reflects the variation of gene expression due to, e.g., cell cycle, sub-cell-type composition, cell-level covariates such as ribosome RNA fraction. The estimation of an NBMM, in general, is computationally intensive, especially for large sample size because the marginal likelihood is intractable due to high-dimensional integrals of the random effects. A standard estimation method for generalized linear mixed models (GLMMs) using penalized quasi-likelihood (PQL) (Breslow and Clayton, 1993) involves a two-step iteration procedure where the variance of the random effects is estimated first, and then it is used to estimate the coefficients of the variables of interest. In genome-wide association studies (GWAS), the variance component is often estimated only once under the null model. In contrast, such a two-step procedure has to be carried out for each of tens of thousands of genes in scRNA-seq data or peak regions in scATAC-seq data, which is considerably more challenging. NEBULA achieves its speed gain by using an approximate analytical marginal likelihood, which was derived leveraging the fact that the data for each subject often include a large number of cells. This approximation, referred to as NEBULA-LN, eliminates the computationally-demanding two-layer high-dimensional optimization procedure, and therefore substantially reduces the computational time for estimating the overdispersions. For situations in which NEBULA-LN fails to achieve accurate estimates of the subject-level overdispersion, we estimate the cell-level overdispersion using NEBULA-LN and implemented a fast estimation procedure, referred to as NEBULA-HL, based on a hierarchical likelihood (h-likelihood) to estimate the subject-level overdispersion. Because only one overdispersion is needed to be optimized in the h-likelihood, NEBULA-HL converges in much fewer steps.

### NEBULA is computationally efficient

We compared NEBULA with the existing statistical tools for estimating NBMMs to evaluate its computational efficiency. In addition to NEBULA-LN, we assessed a combination of NEBULA-LN and NEBULA-HL (denoted by NEBULA-LN+HL), in which we first used NEBULA-LN to estimate the cell-level overdispersion, and then fixed this estimate in NEBULA-HL to obtain the subject-level overdispersion. We chose the L-BFGS algorithm from (Ypma, 2014) for optimization in NEBULA-LN. We found that a Newton-Raphson (NR) algorithm (Dennis and Schnabel, 1996) was generally faster in this case but less robust in terms of convergence. For comparison, we included the *glmer*.*nb* function from the *lme4* R package (Bates et al., 2014), the *glmmTMB* function from the *glmmTMB* R package (Brooks et al., 2017), and the *inla* function from the *INLA* R package (Rue et al., 2009). The *glmmTMB* R package utilizes the Template Model Builder automatic differentiation engine, and the *INLA* R package is an efficient Bayesian framework based on integrated nested Laplace approximations. In *glmer*.*nb*, we assessed the default setting (nAGQ=1), which is based on Laplace approximation (LA) (Tierney and Kadane, 1986), and a faster but less accurate setting (nAGQ=0), which is based on a penalized likelihood (Lindstrom and Bates, 1990). The difference is that LA estimates the fixed effects in the marginal likelihood with the variance components, while the latter estimates the fixed effects with the random effects in the penalized likelihood. We did not include methods using adaptive Gaussian quadrature (AGQ) (*glmer*.*nb* with nAGQ>1) (Pinheiro and Bates, 1995; Pinheiro and Chao, 2006) or Bayesian methods using Markov chain Monte Carlo (MCMC) (Vestal et al., 2020) because of their computational intensity. In addition, the Gaussian variational approximation method (Ormerod and Wand, 2012) is promising, but has not been implemented for the NBMM.

Given a simulated data set comprising 50 subjects and CPS=200 (a total of ∼10,000 cells), it took ∼400s for NEBULA-LN and ∼1000s for NEBULA-LN+HL to analyze 10,000 genes (Fig. 1A). These benchmarks were on average >100-fold and ∼50-fold faster than *glmer*.*nb* with nAGQ=0, respectively. It took much longer for *glmer*.*nb* with the default setting and *glmmTMB*. We failed to obtain the full benchmarks for *inla* because it took an extremely long time to finish in some scenarios. Generally, the running time of *inla* was between *glmmTMB* and the default *glmer*.*nb*. The computational time of all methods increased almost linearly with respect to CPS, which was expected because the random effects are assumed to be independent. We also observed that the computational time of all methods scaled in a similar trend with the increasing number of subjects (Fig. 1B). In all settings, NEBULA-LN+HL was ∼2-3x slower than NEBULA-LN but was still much faster than *glmer*.*nb*. In our following analysis of ∼34,000 excitatory neurons from 48 individuals in the snRNA-seq data adopted from (Mathys et al., 2019), NEBULA accomplished an association analysis of ∼16,000 genes for identifying marker genes in ∼40 minutes, compared to ∼67 hours using *glmer*.*nb* with nAGQ=0. When the number of fixed-effects predictors increased from two to ten, the computational time of both NEBULA and *glmer*.*nb* with nAGQ=0 grew modestly, while the default *glmer*.*nb* increased by ∼10x (Fig. S1). This substantial increase of the computational time in the default *glmer*.*nb* is expected because it estimates the fixed effects with the overdispersions in the outer layer of the iterative procedure using a derivative-free method. The number of iterations of such an optimization method rises drastically when the dimension of the parameter space increases.

**Figure 1.**
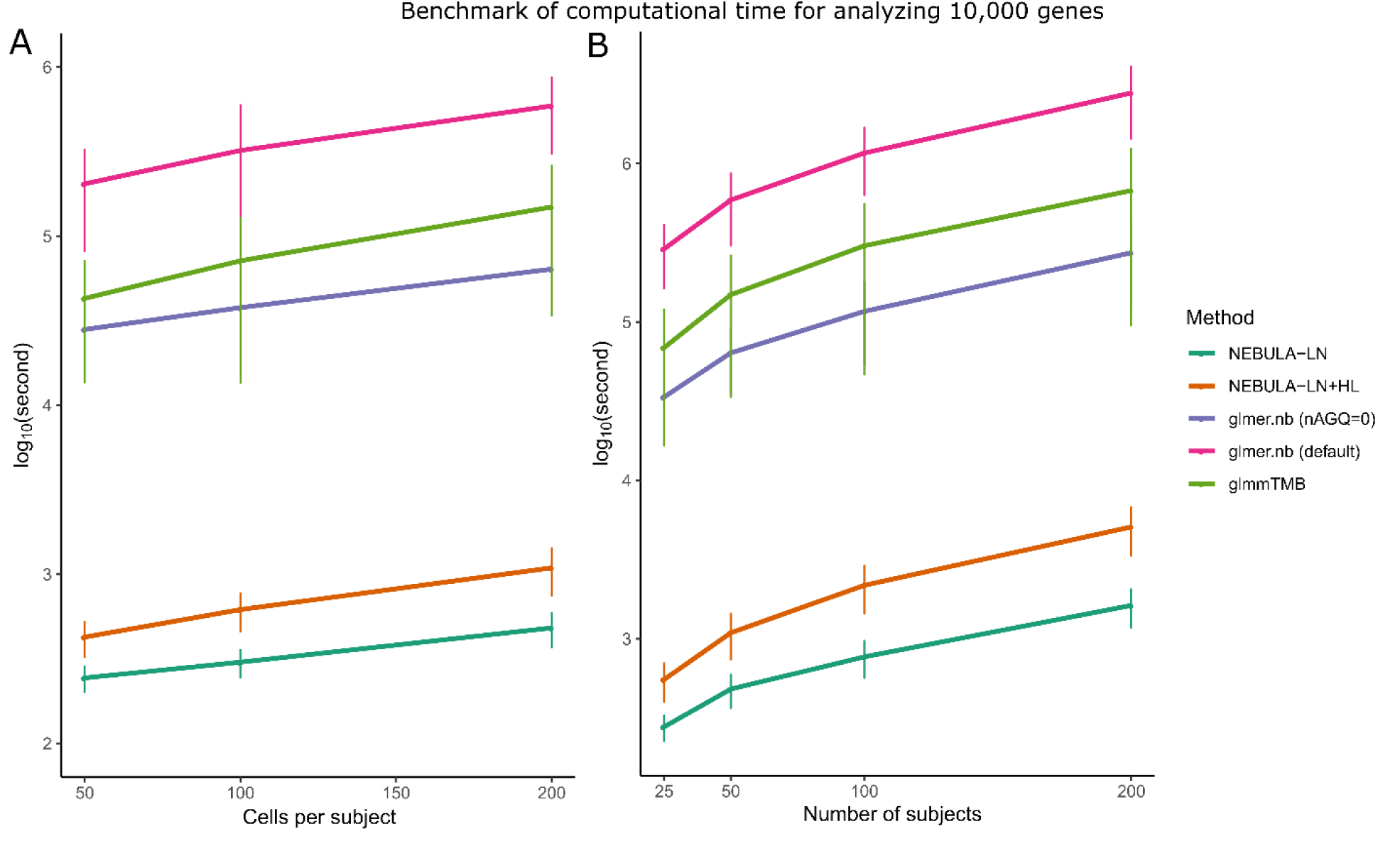
(A) The computational time (measured in log10(seconds)) of fitting an NBMM for 10,000 genes with respect to CPS. The number of subjects was set at 50. (B) The computational time (measured in log10(seconds)) of fitting an NBMM for 10,000 genes with respect to the number of subjects. The CPS value was set at 200. Two fixed-effects predictors were included in the NBMM. The average benchmarks were summarized from scenarios of varying subject-level and cell-level overdispersions and the CPC value of a gene ranging from exp(−4) to 1. CPS: cells per subject. CPC: counts per cell.

### Decomposing overdispersions using NEBULA

We show in Supplementary materials A.5 that the approximation proposed in NEBULA-LN still provides asymptotically consistent estimates. Nevertheless, it is also crucial to assess its practical performance under a finite sample size. As the above benchmarks indicated that NEBULA-LN had a huge speed advantage over NEBULA-HL, the basic strategy in NEBULA is to first fit the data using NEBULA-LN and apply NEBULA-HL only in situations where NEBULA-LN performs poorly. We, therefore, conducted a comprehensive simulation study to investigate under which situations NEBULA-LN could produce sufficiently accurate estimates. The estimation accuracy of NEBULA-LN is primarily affected by CPS, the magnitude of the cell-level dispersion, and the coefficient of variation (CV) of the scaling factor. We thus focus on determining a threshold in terms of these parameters. Because NEBULA-HL was based on a standard h-likelihood method, we used NEBULA-HL as a reference to assess the accuracy of NEBULA-LN in terms of mean square error (MSE).

We first evaluated the performance of NEBULA-LN in a well-designed study where the numbers of cells were highly homogeneous across the subjects and were assumed to follow a Poisson distribution. We considered the CPS value varying from 100 to 800, and the number of subjects ranging from 30 to 100, which are common settings in real droplet-based scRNA-seq data. We simulated counts based on an NBMM with the subject-level overdispersion *σ*^2^ ranging from 0.01 to 1 and the cell-level overdispersion *ϕ*^−1^ ranging from 0.01 to 100 (A larger *ϕ*^−1^ means higher overdispersion). These ranges were selected because they were close to our estimates observed in the real snRNA-seq dataset in (Mathys et al., 2019). We simulated genes of different expression level by tuning *β*_0_ from −4 to 2 (When the subject-level overdispersion *σ*^2^ is small, exp (*β*_0_) approximately equals the counts per cell (CPC) defined by the total counts of a gene divided by the total number of cells). We considered the following two situations, (i) all cells share the same scaling factor and (ii) the scaling factor across cells varied substantially with the coefficient of variation equal to 1.

Fig. 2A shows that the MSE of NEBULA-LN for estimating the cell-level overdispersion *ϕ*^−1^ decreased uniformly with increasing CPS. The convergence of the estimates to their true values supported our theoretical conclusion that NEBULA-LN is asymptotically consistent for estimating *ϕ*. The MSEs of both methods increased when the mean expression of a gene (measured by CPC) dropped. Both methods struggled to accurately estimate *ϕ* for low-expressed genes when CPC≈0.018, and the bias of NEBULA-LN became more noticeable. This suggests that genes with a low CPC value could be filtered during the quality control (QC) because it is difficult to obtain accurate estimates of the overdispersions for these genes due to the high sparsity of the data unless the number of cells is very large. In most scenarios, NEBULA-LN showed higher MSEs and was asymptotically less efficient than NEBULA-HL. Nevertheless, the MSEs between the two methods were comparable when CPS was >400. When *ϕ* was very large (=100), NEBULA-LN produced a more biased estimate if CPS is small. In addition, Fig. S3 shows that increasing the number of subjects alone improved the standard error, but not the bias of 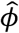, suggesting that the bias of 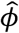 depended mainly on CPS. A larger CV of the scaling factor showed little impact on the MSE of NEBULA-LN (Fig. S2A).

**Figure 2.**
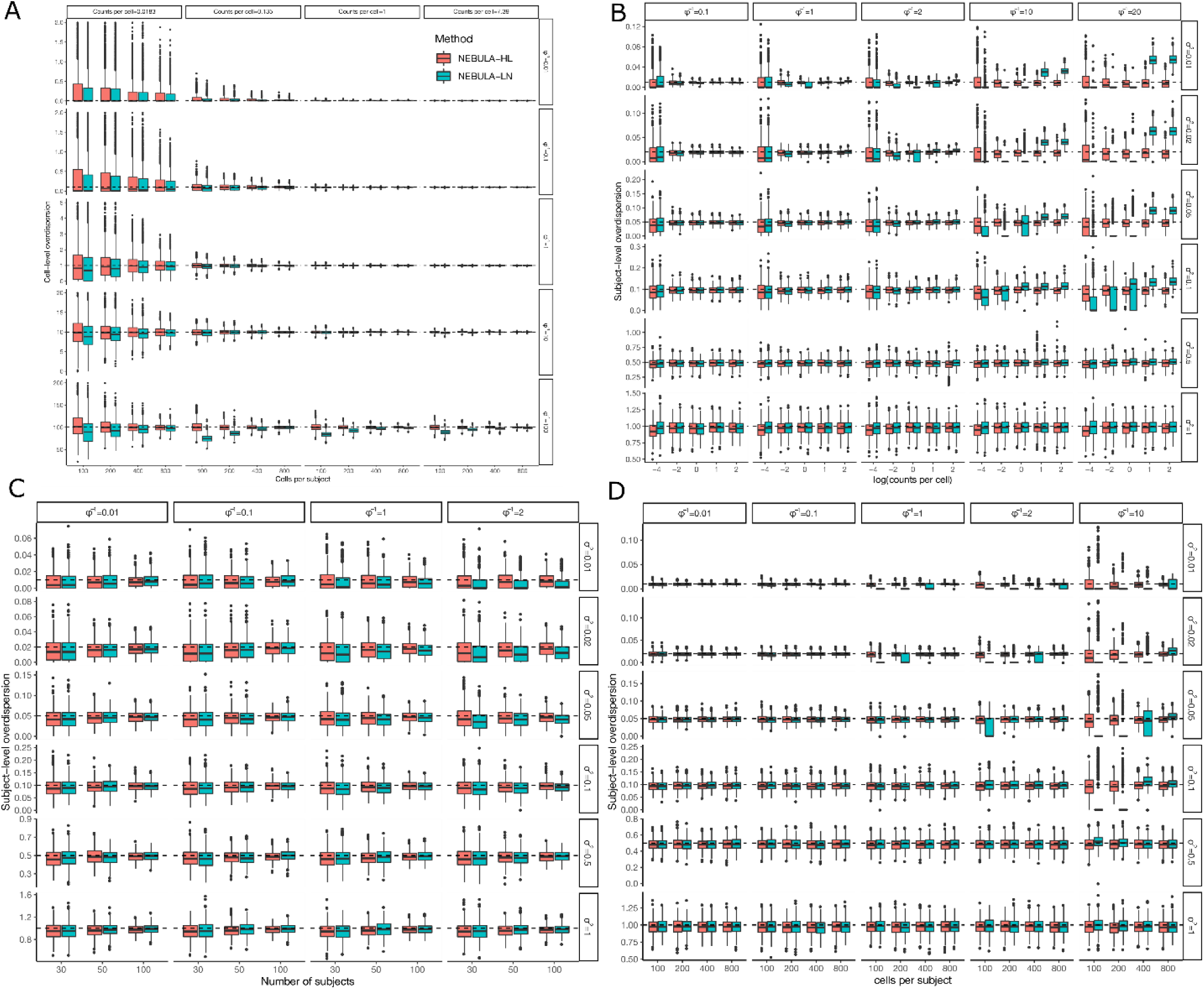
Comparison of estimated cell-level and subject-level overdispersions between NEBULA-LN and NEBULA-HL. The summary statistics were calculated from five hundred simulated replicates in each of the scenarios. (A) The cell-level overdispersions estimated by NEBULA-LN and NEBULA-HL under different combinations of CPS, *β*_0_ (a lower *β*_0_ corresponding to a lower CPC value), and *ϕ*. The number of subjects was set at 50. (B) The subject-level overdispersions estimated by NEBULA-LN and NEBULA-HL under different combinations of *β*_0_, *σ*^2^, and *ϕ*. The number of subjects was set at 50. The CPS value was set at 400. (C) The subject-level overdispersions estimated by NEBULA-LN and NEBULA-HL under different combinations of the number of subjects, *σ*^2^, and *ϕ*. The CPS value was set at 400, and *β*_0_ was set at 0.05. (D) The subject-level overdispersions estimated by NEBULA-LN and NEBULA-HL under different combinations of CPS, *σ*^2^, and *ϕ*. The number of individuals was set at 50, and *β*_0_ was set at 1. CPS: cells per subject. CPC: counts per cell.

On the other hand, NEBULA-LN had similar or lower MSEs than NEBULA-HL for estimating the subject-level overdispersion when *σ*^2^ is large (≥0.5) (Fig. 2B). However, when *σ*^2^ was small, NEBULA-LN began to produce CPC-dependent biased estimates when *ϕ* turned larger (Fig. 2B). Increasing CPS alleviated the bias rapidly (Fig. 2D). When the CV of the scaling factor increased to one, an approximately 2-fold increase of CPS · *ϕ* was required to achieve similar MSEs by comparing Figs. S2B, S2D with Figs. 2B, 2D. Combining these observations suggested that the accuracy of NEBULA-LN for estimating *σ*^2^ approximately depends on *κ* = CPS · *ϕ*/(1 + *c*^2^) (*c* is the CV of the scaling factor), which is derived in the Methods section. These empirical results suggested that *κ* determined the lower bound of *σ*^2^ for which an accurate estimate could be achieved. NEBULA-LN was able to obtain good estimates for *σ*^2^ as low as 0.02 when *κ* = 200 (corresponding to *ϕ*^−1^ = 2 in Fig. 2B and Fig. 2C), which was sufficient to reliably test cell-level predictors, as we will show in the next section. Both NEBULA-LN and NEBULA-HL underestimated *σ*^2^ particularly when the number of subjects was small (Fig. 2C). We found that the underestimate was not specific to NEBULA, but also present in other tools based on PQL (e.g., the *lme4* R package with nAGQ=0). Increasing the number of subjects alleviated this bias, and the influence of such bias on testing subject-level preditors will be discussed in the next section.

We further evaluated the performance of NEBULA in unbalanced samples, where the numbers of cells varied substantially across the individuals. An unbalanced sample is common when analyzing a cell subpopulation obtained from clustering. For example, in the snRNA-seq data in (Mathys et al., 2019), several individuals had only ∼20 inhibitory neurons, although the CPS value among the 48 individuals was ∼200. Overall, the results from the unbalanced (Fig. S17) were comparable to those from the balanced sample (Fig. 2), suggesting the distribution of the number of cells had little impact on estimating *σ*^2^ and *ϕ*.

### NEBULA controls type I error rate

As shown above, either NEBULA-LN or NEBULA-HL might yield somewhat biased estimates of the overdispersions in specific scenarios. We then assessed the impact of such biases on controlling the type I error rate by testing the fixed-effects predictors under a wide range of settings. As mentioned above, predictors can be classified as either a cell-level or subject-level variable. For a cell-level predictor, Fig. 3A and Fig. 3B show that both methods controlled the type I error rate well in almost all scenarios. The type I error rate was only slightly inflated when both overdispersions were very large, and the CPS value was as small as 100 (i.e., the bottom-right panels in Fig. 3A and Fig. 3B). Although *ϕ*^−1^ was underestimated in NEBULA-LN more for low-expressed genes than high-expressed genes (Fig. 2A), no inflation of the type I error rate was observed for low-expressed genes (Fig. S7). Besides, biased estimates of the subject-level overdispersion in NEBULA-LN had little effects on testing a cell-level predictor.

**Figure 3.**
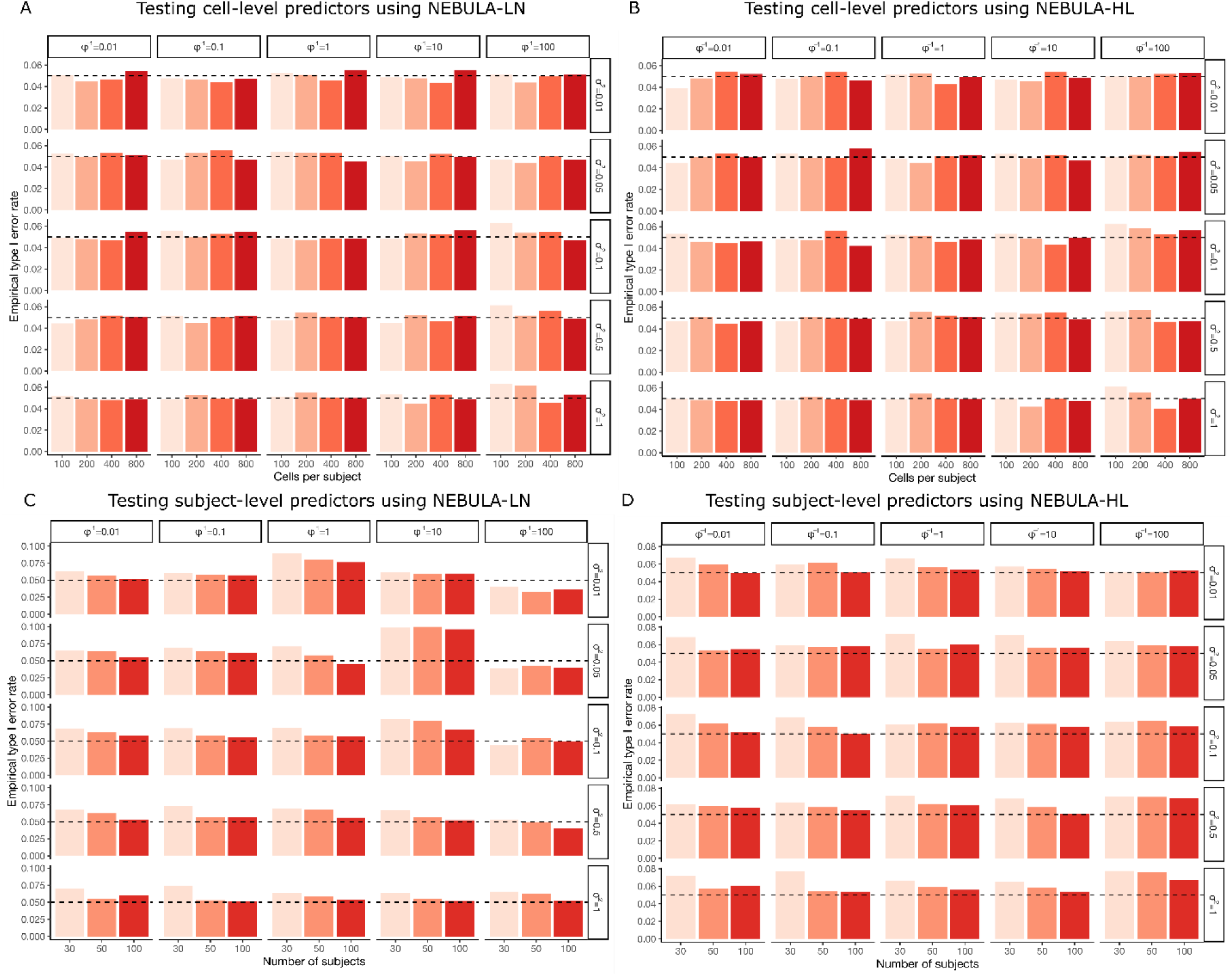
Empirical type I error rate of testing cell-level and subject-level predictors using NEBULA-LN and NEBULA-HL. The type I error rate was calculated from five hundred simulated replicates in each of the scenarios and was evaluated at the significance level of 0.05 (the dashed lines). (A) The empirical type I error rate of NEBULA-LN under different combinations of CPS, *σ*^2^, and *ϕ*. The number of individuals was set at 50. (B) The empirical type I error rate of NEBULA-HL under different combinations of CPS, *σ*^2^, and *ϕ*. The number of subjects was set at 50. (C) The empirical type I error rate of NEBULA-LN under different combinations of the number of subjects, *σ*^2^, and *ϕ*. The CPS value was set at 400. (D) The empirical type I error rate of NEBULA-HL under different combinations of the number of subjects, *σ*^2^, and *ϕ*. The CPS value was set at 400. CPS: cells per subject. CPC: counts per cell.

In contrast, the type I error rate of testing a subject-level predictor was highly sensitive to a biased estimate of the subject-level overdispersion *σ*^2^. The type I error rate in both methods was inflated in the case of 30 subjects and the inflation diminished with the increasing number of subjects (Fig. 3C, Fig. 3D). This is because both methods underestimated the subject-level overdispersion when the number of individuals was small (Fig. 2C). The comparison between Fig. 2C and Fig. 3C suggested that even a small underestimated subject-level overdispersion would lead to an inflated type I error rate of testing a subject-level predictor when the number of subjects is small. Combining Fig. 2B and Fig. S6 suggested that NEBULA-LN had inflated or deflated type I error rate of testing a subject-level predictor whenever *σ*^2^ was overestimated or underestimated, respectively, even for *σ*^2^ being as low as 0.01. Underestimates of the subject-level overdispersion were not specific to NEBULA but also observed in *glmer*.*nb* with a penalized likelihood (nAGQ=0). These results suggested that NEBULA or other estimation methods based on a penalized likelihood should be used with caution when testing a subject-level predictor, particularly when the number of subjects is small (<100).

We also evaluated the performance of a Poisson mixed model for testing a subject-level predictor. If we ignore the cell-level overdispersion *ϕ*^−1^ in NEBULA, we end up with a Poisson gamma mixed model (PGMM) (Sutradhar and Qu, 1998), the likelihood function of which has a simple analytical form. Therefore, it does not suffer from the problem of underestimating *σ*^2^ as in NEBULA-HL and is computationally much faster than the Poisson mixed model estimated by *glmer*.*nb* with nAGQ=10 adopted in (Mathys et al., 2019). Because the cell-level overdispersion is ignored in the PGMM, it definitely cannot be used to test a cell-level predictor. However, it is still interesting to check its type I error rate when used for testing a subject-level predictor. Fig. S15 shows that overall the PGMM controlled the type I error rate well in scenarios where the number of subjects was 100, except for situations where both the cell-level and subject-level overdispersions were large (*ϕ*^−1^ ≥ 100 and *σ*^2^ ≥ 0.5). But such situations account for a very small fraction of genes (usually low-expressed genes) in real data which are often excluded during quality control. For instance, we observed <0.1% genes with CPC>0.1% having *ϕ*^−1^ > 100 in the major cell types in the snRNA-seq data in (Mathys et al., 2019), and the percentage became almost zero when we filtered out low-expressed genes (CPC<1%) during quality control. Therefore, the PGMM might be considered as a fast tool for testing subject-level predictors followed by a second-round sensitivity analysis for top signals if the computational intensity is a major concern.

Based on these simulation results, the strategy that we recommend for testing a cell-level predictor, as summarized in Table S3, is to fit the data first using NEBULA-LN to obtain estimates 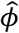 and 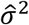. Both overdispersion parameters are re-estimated using NEBULA-HL if 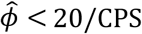 because biased estimates of both overdispersions might occur in this case. Otherwise, we fix 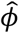 and only use NEBULA-HL to estimate *σ*^2^ if 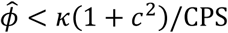, where we chose *κ* = 200 based on the simulation results. We report the estimates directly from NEBULA-LN if 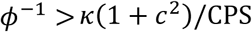 because in this case, NEBULA-LN can produce a reliable estimate for a sufficiently small *σ*^2^. We used this strategy in our following analysis of the snRNA-seq data in the human frontal cortex (Mathys et al., 2019), and adopted the cell library size (the total number of reads) as the scaling factor. Under this strategy, ∼90% and ∼80% of genes with CPC>0.1% were analyzed using NEBULA-LN alone in the excitatory neurons (CPS=∼700) and oligodendrocytes (CPS=∼380). We found that *c*^2^ was ∼0.8 for the neurons and ∼0.25 for all other cell types. The larger CV of the cell library size in the neurons was probably because the neurons consisted of highly heterogeneous subpopulations (Fig. S8). On the other hand, for testing a subject-level predictor, we recommend using *κ* = 1000 or even larger.

### Subpopulation heterogeneity and known covariates contribute to overdispersions

The subject-level and cell-level overdispersion parameters likely reflect some intrinsic biological features of a gene. A gene with a larger subject-level overdispersion might indicate that the gene is strongly regulated by subject-level factors such as age, sex, disease status, exposure, and genetic variation. We then dissected the overdispersions using NEBULA for genes in the snRNA-seq data in (Mathys et al., 2019). We carried out the analysis in each of the six major cell types, including excitatory neurons, inhibitory neurons, oligodendrocytes, astrocytes, oligodendrocyte progenitor cells (OPCs), and microglia.

We observed that the mean cell-level overdispersion grew steadily with lower expression of a gene in the excitatory neurons (Fig. 4A) and the other cell types (Fig. S11). This mean-overdispersion trend is reminiscent of that observed in bulk RNA-seq data, in which low-expressed genes often show higher variance (Law et al., 2014). Less than 0.1% of genes in the excitatory neurons had an estimated cell-level overdispersion 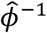 larger than 100, and all of them were low-expressed genes. In contrast, the subject-level overdispersion exhibited a similar mean-overdispersion trend, except that the overdispersion leveled off for abundantly expressed genes in the case of CPC>1 in the excitatory neurons (Fig. 4B). Similar patterns were observed in the other cell types (Fig. S12). We observed a higher variation for both estimated overdispersion in the lower-expressed genes due to the larger standard error for those genes, as shown in the simulation (Fig. 2). There was no clear correlation between the subject-level overdispersion and the cell-level overdispersion (Fig. 4C).

**Figure 4.**
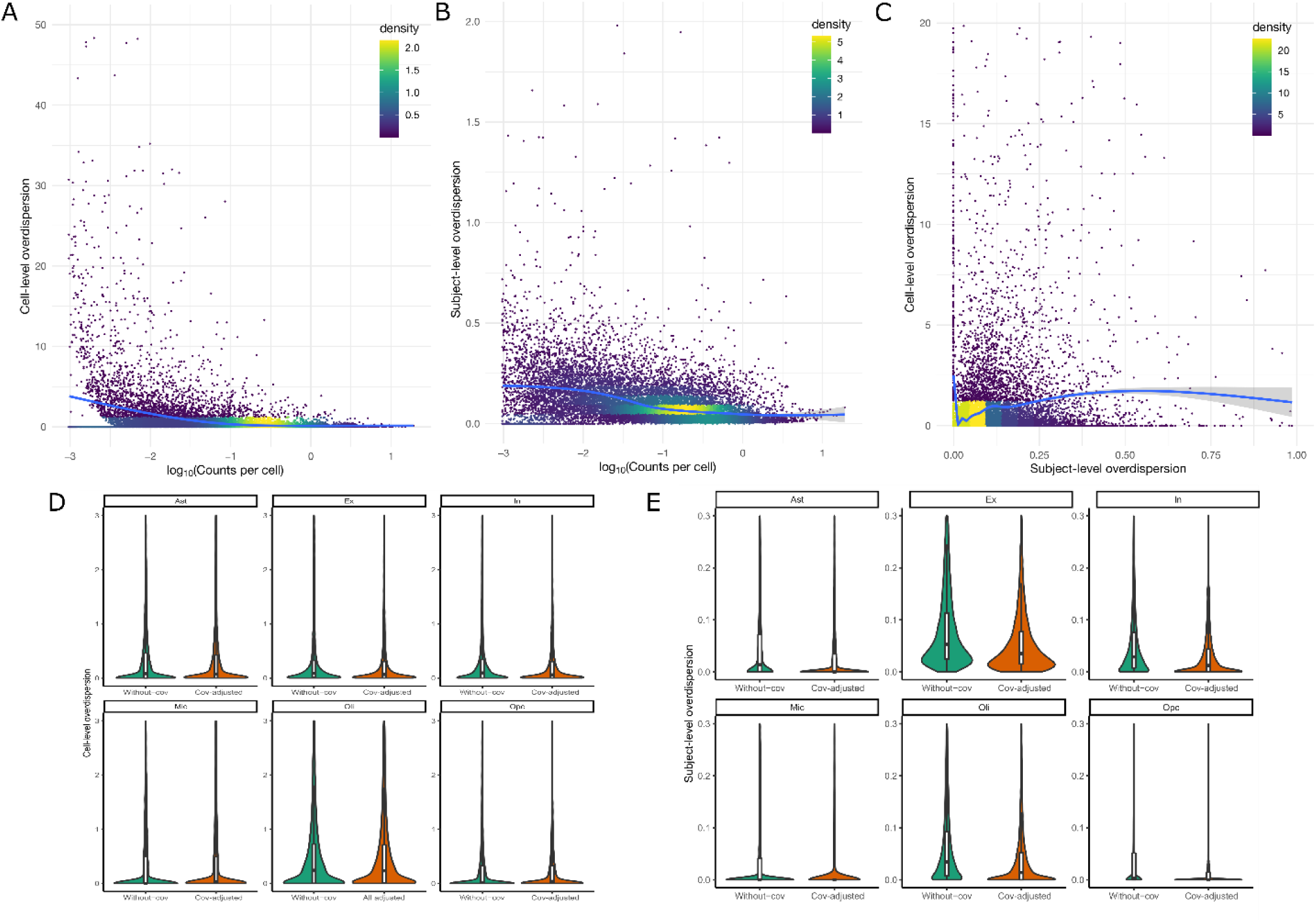
(A) The cell-level overdispersions of 16,207 genes with CPC>0.1% in the excitatory neurons estimated by NEBULA with respect to the CPC of the gene. No covariates other than the intercept were included in the model. The density is the two-dimensional kernel density computed using the *kde2d* R function with n=100. (B) The subject-level overdispersions of 16,207 genes with CPC>0.1% in the excitatory neurons estimated by NEBULA with respect to the CPC of the gene. (C) The cell-level overdispersions of 16,207 genes with CPC>0.1% in the excitatory neurons estimated by NEBULA versus their subject-level overdispersions. (D) The estimated cell-level overdispersions of all genes with CPC>0.1% in each of the six brain cell types before and after adjusting for age, sex, race, AD status, the total number of features of the cell, and the percentage of ribosomal genes. (E) The estimated subject-level overdispersions of all genes with CPC>0.1% in each of the six brain cell types before and after adjusting for age, sex, race, AD status, the total number of features of the cell, and the percentage of ribosomal genes.

We then explored which factors might contribute to both overdispersions of a gene in each of the cell types. First, we found that the median cell-level overdispersion dropped markedly in the subpopulations (Fig. S13A), suggesting that sub-cell-type differential expression was the major source for the cell-level overdispersion. On the other hand, the median subject-level overdispersion dropped to a much smaller extent (Fig. S13B). We then investigated subject-level covariates, including age, sex, AD diagnosis, and race, and cell-level covariates, including the number of total features of a cell and percentage of ribosomal protein genes by adding them to the regression model. We found that the overall subject-level overdispersion dropped substantially in most of the cell types after adjustment for these covariates, but the cell-level overdispersion almost remained at the same level (Fig. 4D, Fig. 4E). These results suggested that the subject-level overdispersion was largely attributed to these factors, while the cell-level overdispersion resulted from subpopulation heterogeneity and other unknown factors, e.g., drop-out rates.

To compare different NBMMs, we applied *glmer*.*nb* in the *lme4* R package to the same data. The estimated cell-level overdispersions were highly comparable between both methods (Fig. S14), except for an outlier in *glmer*.*nb*. The estimated subject-level overdispersions were also highly correlated for those genes with a small to moderate subject-level overdispersion (Fig. S9), although it is not straightforward to compare the absolute values because the random effects in these two NBMMs are assumed to follow slightly different distributions (See the Methods section). The larger deviation between NEBULA and *glmer*.*nb* observed for higher subject-level overdispersion was expected, as previously shown that the two models are almost the same when the subject-level overdispersion is small (Sutradhar and Qu, 1998). Estimated fixed effects were also consistent between NEBULA and *glmer*.*nb* (Fig. S4).

### Identifying sub-cell-type marker genes using NEBULA

Using a negative binomial regression without the subject-level random effects could result in substantially inflated false positives (Milanzi et al., 2012). Disproportionate cell frequency among individuals may exacerbate the inflation of spurious associations, similar to Simpson’s Paradox (Simpson, 1951). In this case, individuals can be a confounding factor between gene expression and cell clusters. On the other hand, adding individuals as fixed effects would lead to too many parameters in the model, especially when the number of individuals is large. Therefore, an NBMM is an ideal approach for the identification of marker genes for subpopulations in multi-subject scRNA-seq data.

For the demonstration, we used the 3386 astrocytes from the snRNA-seq data in the frontal cortex, which were classified into four subclusters (AST0 - AST3). Some of the subclusters were distributed across the subjects in a highly unbalanced pattern. For example, the vast majority of the cells in the AST3 cluster came from only four subjects (Fig. S10). We simulated the counts of gene expression using an NBMM under the null hypothesis (i.e., not associated with any of the subclusters) with both subject-level and cell-level overdispersions. We assessed the false-positive rates (FPR) under four scenarios with the variance component of the subject-level overdispersion ranging from 0 to 0.1. As we can see in Fig. 5A, even with the subject-level overdispersion *σ*^2^ as small as 0.05, the FPR started to show inflation. The inflation rose rapidly with the increased subject-level overdispersion. In contrast, NEBULA controlled the FPR very well.

**Figure 5.**
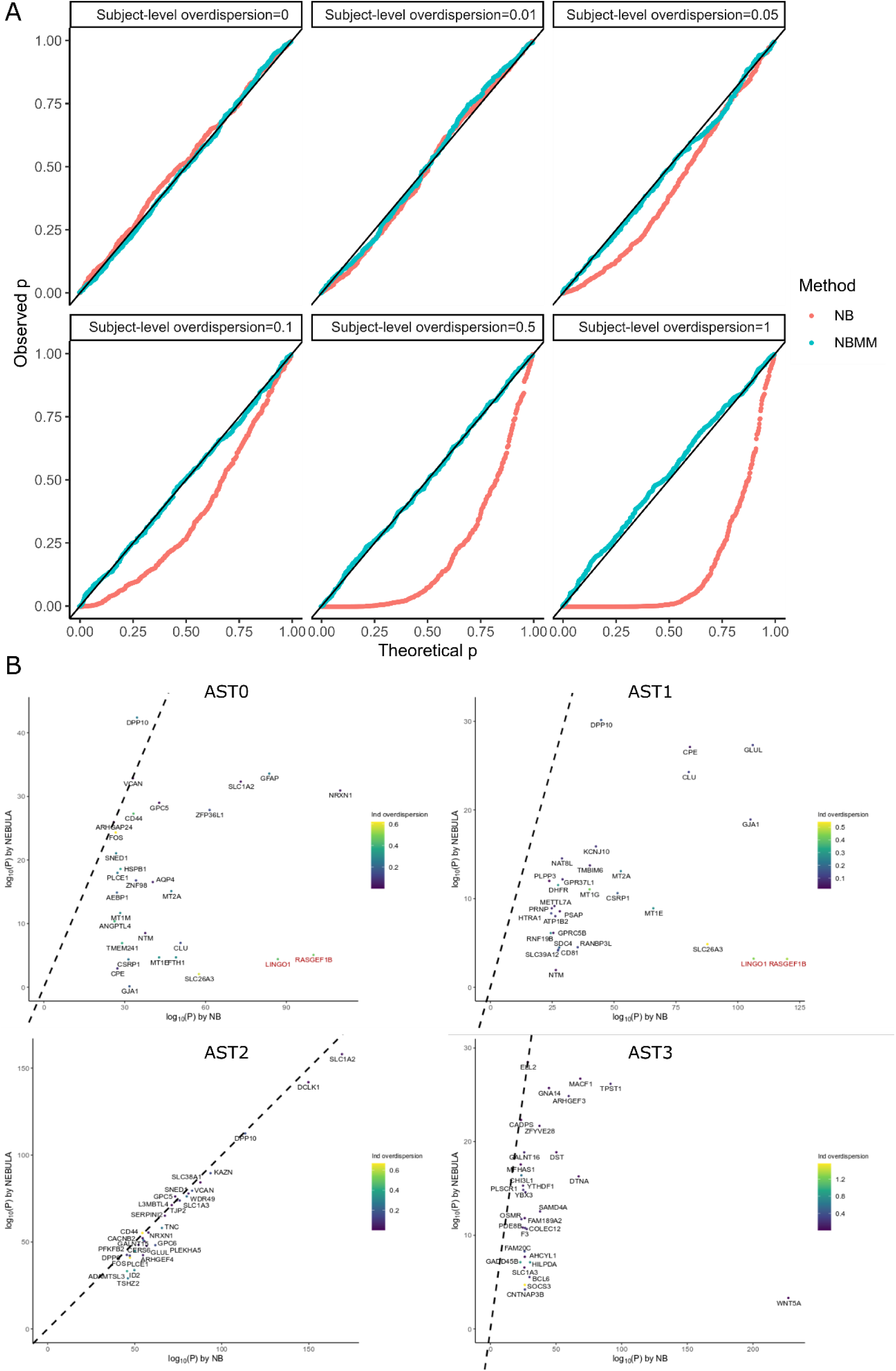
(A) Q-Q plots of p-values for testing the association between simulated counts and cell clusters using NEBULA and a negative binomial regression. The counts were simulated under the null model (i.e., no association between the counts and cell clusters) based on an NBMM with the subject-level overdispersion (*σ*^2^) ranging from 0 to 1 and a cell-level overdispersion fixed at 1. (B) Comparison of the p-values of the top 30 genes identified by a negative binomial regression associated with each of the four astrocyte subpopulations (AST0-AST3) in the snRNA-seq data (Mathys et al., 2019). The labels of the four astrocyte subpopulations are corresponding to those presented in (Mathys et al., 2019). The color of each point gives the subject-level overdispersion of this gene estimated by NEBULA. Two genes (*RASGEF1B* and *LINGO1*) highlighted in red showed highly reduced significance in NEBULA compared to the p-values in the negative binomial regression.

To evaluate how this inflation of FPR might affect the selection of marker genes, we carried out association analysis between the gene expression and the four subclusters in the astrocytes in the snRNA-seq data (Mathys et al., 2019) using a negative binomial regression and NEBULA. Among the top 30 genes identified by a negative binomial regression, most of the p-values were consistent between the two methods in the subclusters AST2 and AST3. However, some of the top genes in the subclusters AST0 and AST1, such as *RASGEF1B* and *LINGO1*, were much less significant in NEBULA than in the negative binomial regression (Fig. 5B). Both genes showed uniformly strong subject-level overdispersion in the astrocytes (Fig. 5B) and the other five cell types, indicating that the highly significant p-values in the negative binomial regression could result from failing to account for the subject-level overdispersion. Furthermore, an independent cell-type-specific bulk RNA-seq data (Zhang et al., 2016) showed that the expression of *RASGEF1B* and *LINGO1* was much lower in astrocytes than that in microglia and oligodendrocytes, respectively. Given the evidence, it should be cautious about classifying these genes as subcell-type marker genes based on the results from the negative binomial regression without the random effects. Besides, *GJA1* was identified by NEBULA as a marker gene of AST1 but not AST0, while a negative binomial regression identified *GJA1* as significant in both subclusters (Fig. 5B).

### Cell-level co-expression analysis of *APOE*

Estimating co-expression between genes is crucial for building interaction or regulatory networks and detecting biological pathways. However, co-expression estimated from bulk RNA-seq data is heavily affected by many hidden confounders such as cell type composition and experimental noise. In contrast, scRNA-seq data opened up new possibilities for investigating co-expression at the cellular level. We interpret the cell-level co-expression as a co-occurrence of two genes in a cell. Because cells with larger sequencing depth or lower ribosomal protein gene expression had higher co-occurrence rate, we included the library size as the normalizing factor, and the number of features and percentage of ribosomal protein genes as the covariates. As *APOE* was abundantly expressed in astrocytes and microglia in the frontal cortex, we identified genes co-expressed with *APOE* in these two cell types by utilizing NEBULA. We normalized the raw *APOE* UMI counts by the library size of each cell and subtracted the mean *APOE* expression of each individual so that the centered normalized *APOE* expression did not correlate with subject-level covariates.

We observed that *APOE* expression in astrocytes was most correlated with *CLU*, one of the top genes from AD GWAS (Harold et al., 2009), on top of other AD-related genes such as *CST3* (Deng et al., 2001) (Table 1). In microglia, *APOE* was co-expressed with multiple immune-related genes, particularly *TREM2* and those involved in the complement system including *C1QA, C1QB*, and *C1QC* (Table 1). *ITM2B*, an inhibitor of the amyloid-beta peptide aggregation, was strongly co-expressed with *APOE* in both cell types. It should be noted that the identified co-expression was based on the within-subject expression, thus ruled out effects from subject-level confounders such as AD status. We then carried out an isoform-specific co-expression analysis by dividing cells into *APOE e3e3, APOE e4*^+^, and *APOE e2e3*. The co-expression patterns showed substantial differences among the isoform groups in microglia, but not in astrocytes (Table S1). More specifically, strong co-expression was observed in the *APOE e2e3* cell population for *CST3*, and the three genes (*C1QA, C1QB, C1QC*) in the complement system, but not for *TREM2* (Table S2), which might suggest different biological mechanisms in microglia behind these two *APOE* isoforms.

**Table 1:**
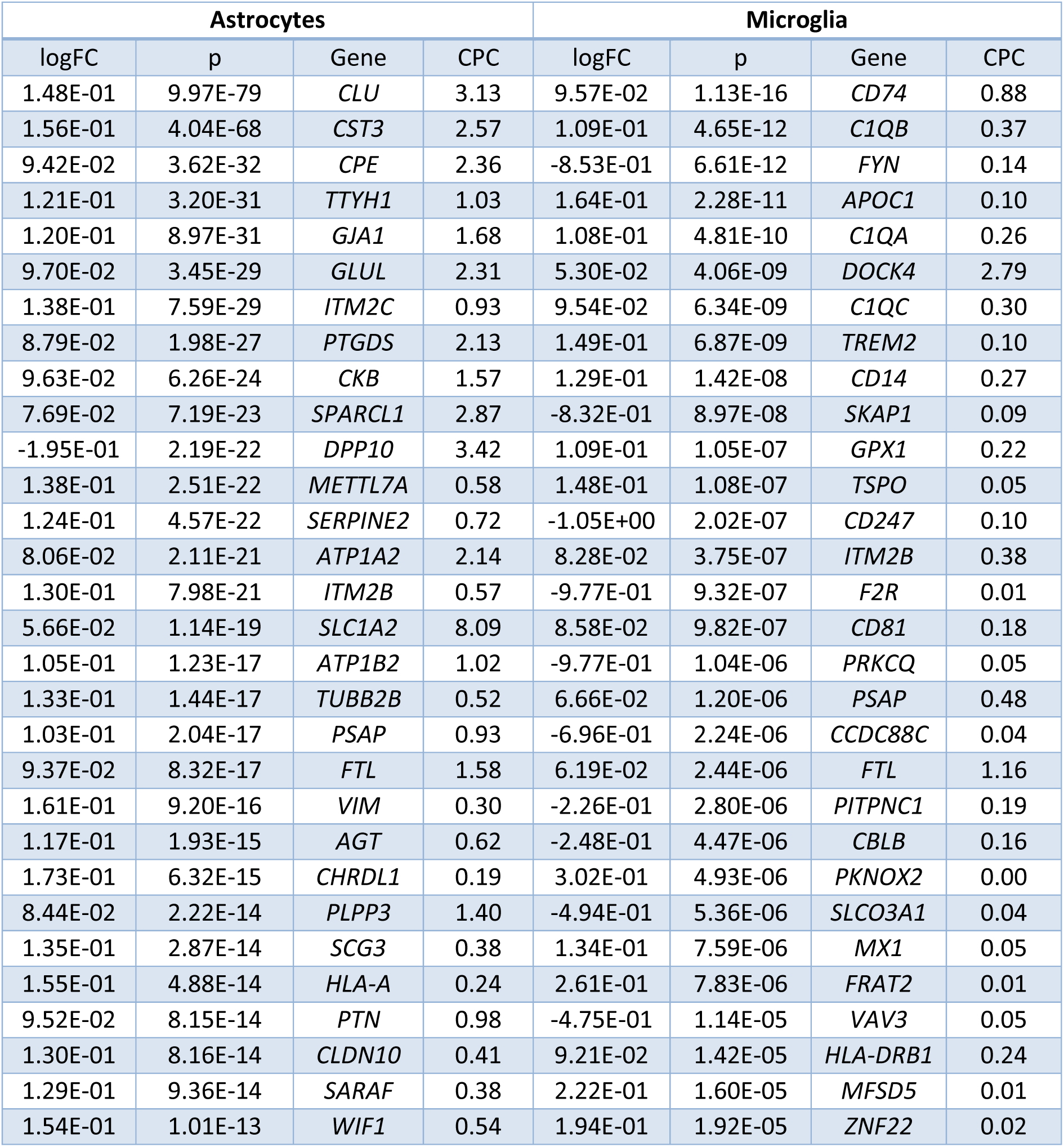
Top 30 genes co-expressed with *APOE* at the single-cell level in astrocytes and microglia in the human frontal cortex. logFC: The log(fold change) of the gene expression if the *APOE* expression is increased. logFC>0 means a positive correlation, and logFC<0 means a negative correlation with the expression of *APOE*. CPC: The UMI counts per cell of this gene in the specific cell type.

## Discussion

In this work, we propose a fast NBMM for the association analysis of large-scale multi-subject single-cell data. We evaluated and demonstrated the performance of NEBULA using a comprehensive simulation study and a droplet-based snRNA-seq data set in the human frontal cortex. Overall, our results show that combining methods based on the approximated marginal likelihood and the h-likelihood, NEBULA managed to achieve significant speed gain and also practically preserve estimation accuracy for analyzing scRNA-seq data. We expect that NEBULA can also be appropriate for other single-cell data such as scATAC-seq data.

The asymptotic efficiency and estimation accuracy of NEBULA-LN are primarily determined by CPS and the cell-level overdispersion. Specifically, our results suggest that *ϕ* · CPS > 20 was sufficient to produce a highly accurate estimate of the cell-level overdispersion. This means that genes with a cell-level overdispersion <10 in a single-cell data set with ∼200 cells per subject can benefit from NEBULA-LN. This criterion covers the vast majority of genes in many large-scale real data. On the other hand, a larger CPS value enables a more accurate estimate for a smaller subject-level overdispersion. Accurate estimation requires a larger *ϕ* · CPS for the subject-level overdispersion than the cell-level overdispersion and also depends on the CV of the scaling factor. Compared to a standard h-likelihood method, we see that NEBULA-LN sacrifices some asymptotic efficiency in estimating the cell-level overdispersion, which means that more cells and counts per cell are needed to achieve the same MSEs. This is the price to pay for the gain of computational efficiency.

NEBULA is solid in controlling the FPR for testing cell-level variables such as subpopulation marker genes. The simulation study suggested that although the estimated overdispersions for low-expressed genes were often less accurate, there was little influence on the FPR. It is possible to treat subjects as batch effects and regress them out in identifying marker genes, but this approach would compromise statistical power if the number of subjects is large. NEBULA is a powerful tool for robustly identifying marker genes. On the other hand, our simulation study suggests that testing a subject-level predictor is sensitive to the estimate of the subject-level overdispersion. The downward bias of NEBULA-HL in estimating the subject-level overdispersion is in line with theoretical results for binary data (Breslow and Lin, 1995) and count responses (Lin, 2007). This bias is more severe in genes with very low expression, probably because the LA works poorly when the response is sparse. Therefore, it should be cautious about using a penalized likelihood method or a first-order LA in an NBMM for investigating a subject-level predictor, particularly when the number of subjects is small. An empirical simulation study indicates that LA may work better for count data than binary data, and the bias decreases as the cluster size increases (Joe, 2008). AGQ can alleviate this issue by producing less biased estimates, but it is computationally more intensive (Joe, 2008; Rabe-Hesketh et al., 2002). If the computational efficiency is a major concern, we found that using the PGMM followed by a sensitivity analysis for top findings could be a fast and viable option for testing a subject-level predictor. Another strategy for testing a subject-level variable is to pool all cells of each subject and treat the data as a bulk RNA-seq dataset. One potential disadvantage of this strategy is that it cannot adjust for cell-level covariates such as mitochondrial and ribosomal gene concentration to increase the statistical power.

The trend of higher average cell-level overdispersion in low-expressed genes is in line with those reported in bulk RNA-seq data (Law et al., 2014; Love et al., 2014). A justification for this observation unique to scRNA-seq data can be that low-expressed genes might have a higher drop-out rate or have larger variation across subpopulations. In bulk RNA-seq data, a large proportion of overdispersion results from heterogeneous cell type composition across the individuals. While this heterogeneity is much better controlled in the scRNA-seq data, subpopulation composition still accounts for most of the cell-level overdispersion. The observed smaller cell-level overdispersion in abundant genes might be attributed to lower drop-out rate and stable expression across subpopulations. In contrast, the subject-level overdispersion is primarily explained by known covariates. With increased gene expression, the cell-level overdispersion drops while the subject-level overdispersion stabilizes. More work is needed to examine to what extent the subject-level overdispersion is explained by the effects of *cis*-eQTLs and whether eGenes show a larger subject-level overdispersion.

The co-expression of *APOE* in astrocytes and microglia is intriguing. Among the top co-expressed genes, multiple SNPs in *CLU* and *TREM2* are AD GWAS loci (Guerreiro et al., 2013; Harold et al., 2009; Jonsson et al., 2013; Lambert et al., 2009). *CLU, CST3*, and *ITM2B* are involved in regulating beta-amyloid production or its fibril formation (Bell et al., 2007; Kaeser et al., 2007; Kim et al., 2008; Matsubara et al., 1995; Matsuda et al., 2005). *CST3* and *ITM2B* are culprits implicated in hereditary cerebral amyloid angiopathy (Revesz et al., 2009). The co-expression results in astrocytes suggest that *APOE* might be involved in the same biological pathway of these genes regulating beta-amyloid production. On the other hand, the co-expression results in microglia support a recent finding (Yin et al., 2019) that *APOE* is involved in the immune system through the complement system, particularly C1q. We further found that this co-expression pattern is *APOE2*^+^ and *APOE4*^+^ dependent in microglia, which might provide more insights into the different role of *APOE* isoforms in AD. These results suggest that the effect of *APOE* on AD is related to regulating both beta-amyloid deposition and the immune system. It should be noted that, because of the data sparsity in the scRNA-seq data, a drawback of this co-expression analysis is that there is little statistical power to identify low-expressed genes because the gene counts have very small variation. Therefore, no observed correlation does not necessarily mean that the two genes are not co-expressed. Interpretation of the co-expression results for low-expressed genes should be cautious.

## Methods

### The model specification in NEBULA

Consider a raw count matrix of *n* cells from *m* individuals in a single-cell data set. We choose to use a negative binomial model to take into account the uncertainty originating from the Poisson sampling process, biological effects, and technical noise. Denote by *y*_*ij*_ the raw count of a gene and by *x*_*ij*_ = (*x*_*ij*0_, …, *x*_*ijk*_) ∈ ℝ^1×(*k*+1)^ a row vector of *k* fixed-effects predictors and the intercept (*x*_*ij*0_ = 1) of cell *j* (*j* ∈ {1, …, *n*_*i*_} and ∑_*i*_ *n*_*i*_ = *n*) in individual *i* (*i* ∈ {1, …, *m*}). We model the raw count using the following NBMM

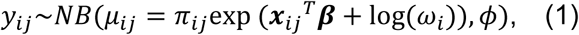

where *NB* is a negative binomial distribution parametrized by its mean *μ* and cell-level dispersion parameter *ϕ* such that the variance is *μ* + *μ*^2^/*φ*, ***β*** ∈ ℝ^(*k*+1)×1^ is a vector of the coefficients of the predictors, *ω*_*i*_ are subject-level random effects capturing the between-subject overdispersion, and *π*_*ij*_ is a known cell-specific scaling factor reflecting the sequencing depth, which can be the library size or some global normalizing factor (e.g., (Robinson and Oshlack, 2010)). Note that *π*_*ij*_ can also be gene-specific as recent studies show that a single normalizing factor might be insufficent to normalize scRNA-seq data (Bacher et al., 2017). The negative binomial log-linear mixed model (NBLMM) proposed in (Booth et al., 2003) assumes a log-normal distribution for the random effects *ω*_*i*_, i.e.,

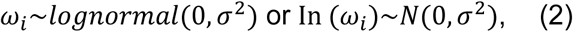

where *σ*^2^ is the variance component of the between-subject random effects.

The estimation of the NBLMM poses a computational challenge because the marginal likelihood is intractable after integrating out the random effects *ω*_*i*_. An estimation method based on an Expectation-Maximization (EM) algorithm is proposed in (Booth et al., 2003). As the NBLMM falls in the framework of the GLMM, conventional estimation methods for the GLMM can be applied, including penalized likelihood-based methods (Breslow and Clayton, 1993; Lindstrom and Bates, 1990), Laplace approximation (Tierney and Kadane, 1986), AGQ (Pinheiro and Bates, 1995; Pinheiro and Chao, 2006). These methods, implemented in the *lme4* R package (Bates et al., 2014) through the argument *nAGQ*, differ in how to approximate the high-dimensional integral in the marginal likelihood, among which the penalized likelihood-based method is the fastest but the least accurate while AGQ is more accurate and much slower. We refer to (Tuerlinckx et al., 2006) for more extensive reviews of these methods. More recently, (Zhang et al., 2017) proposes an estimation procedure for the NBLMM by transforming the problem into repeatedly fitting a linear mixed model using iteratively reweighted least square. All these methods involve at least a two-layer optimization procedure. Although the time complexity is 𝒪(*n*) under the assumption of the independent random effects, the two-layer procedure often requires hundreds of iterations to converge, which becomes the major computational bottleneck. On the other hand, Bayesian methods such as MCMC (Vestal et al., 2020) offer better accuracy, but are even slower than the above methods.

The rationale behind NEBULA is to skip the time-consuming two-layer optimization algorithm for as many genes as possible and to significantly reduce the number of iterations if such an algorithm has to be applied. The key idea is to derive an approximate marginal likelihood in a closed-form so that the model can be estimated using a simple NR or quasi-Newton method. Inspired by (Sutradhar and Qu, 1998), instead of the log-normal distribution in eq. (2), we assume that the between-subject random effects follow the following gamma distribution

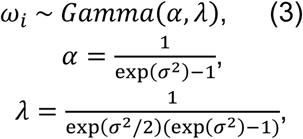

where *α* and *λ* are the shape and rate parameters, respectively. One advantage of this parametrization is that it matches the first two moments to the NBLMM, so that even in a situation where eq. (2) is the true distribution of the random effects, eq. (3) can still provide an accurate estimate of *σ*^2^ when it is not large (Sutradhar and Qu, 1998). Previous studies (Milanzi et al., 2012; Neuhaus and McCulloch, 2011; Neuhaus et al., 2013) indicate that it has practically little influence on the estimation of the fixed-effects predictors to assume a lognormal, gamma, or even misspecified distributions for the random effects, which is also confirmed in our real data analysis (Fig. S4). We call the NBMM with the random effects in eq. (3) as a negative binomial gamma mixed model (NBGMM). Under the NBGMM, the marginal mean and variance of *y*_*ij*_ are given by

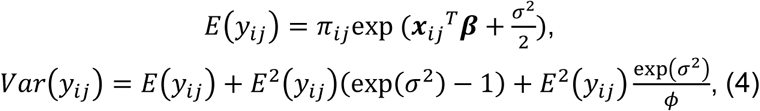

from which we see the first overdispersion term in eq. (4) is controlled by the between-subject variance component *σ*^2^ alone, and the other includes both *σ*^2^ and *ϕ* (The derivation can be found in Supplementary materials A.1). In general, the marginal likelihood of the NBGMM after integrating out *ω*_*i*_ cannot be obtained analytically. However, we show in the next section how to achieve an approximated closed form of the marginal likelihood when *n*_*i*_ is large. We first focus on the derivation in an asymptotic situation, and then investigate its performance in a finite sample.

### Approximation of marginal likelihood in NEBULA-LN

We switch the order of the integral in the marginal likelihood to first integrate out *ω*_*i*_ by rewriting the NBGMM in terms of a mixture of Poisson distributions as follows

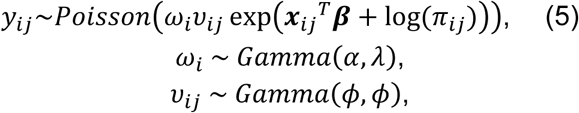

where *v*_*ij*_ has the mixing gamma distribution. The PGMM proposed in (Sutradhar and Qu, 1998), which has a closed-form likelihood, is a special case of the NBGMM if we assume *v*_*ij*_ = 1 in eq. (5). As *ω*_*i*_ is a conjugate random effect for the Poisson model (Molenberghs et al., 2010), a partial marginal likelihood after integrating out *ω*_*i*_ for cells in individual*i* can be expressed explicitly as

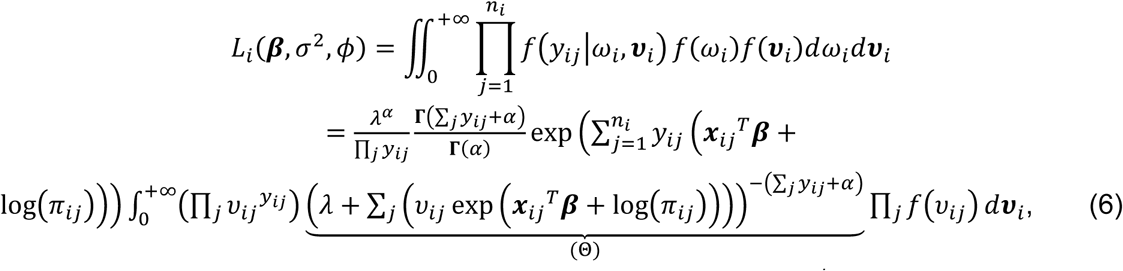

where **Γ**(·) is the gamma function, 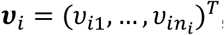, and 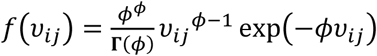 is the gamma density function defined in the NBGMM. Generally, it is not straightforward to solve the high-dimensional integral in eq. (6) because of the entanglement of *v*_*ij*_ in Θ, but when *n*_*i*_ → ∞, we expect to obtain a simple approximation of the summation in Θ asymptotically by the law of large numbers (LLN). Keeping this notion in mind, we rewrite Θ in eq. (6) as

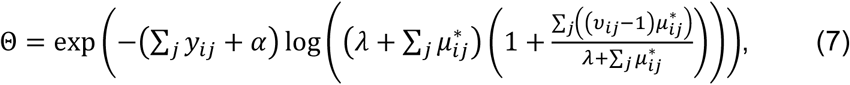

where 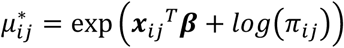. Now consider a Monte Carlo method to approximate the integral in eq. (6) by sampling a large number of ***v***_*i*_ from *f*(***v***_*i*_) and summing up the integrand evaluated at these ***v***_*i*_. We show in Supplementary materials A.2 that under very mild conditions, given any 0 < ε ≪ 1, we have 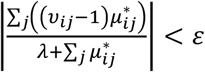 for almost all Monte Carlo samples when *n*_i_ → ∞, that is, 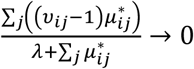 in probability. Given this observation, applying a first-order Taylor expansion to the logarithm in eq. (7), we can approximate the integral in eq. (6) (See Supplementary materials A.3 for the derivation) by

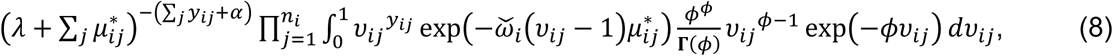

where 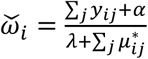. The integrand in eq. (8) is now recognized as a kernel of a gamma distribution with respective to *v*_*ij*_. Calculating this integral explicitly and substituting it into eq. (6) gives an approximated marginal log-likelihood

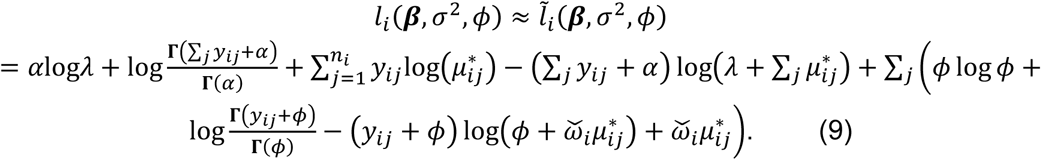

Thus, we may estimate (***β***, *σ*^2^, *ϕ*) by optimizing 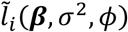, leading to an M-estimator (Huber, 1964, 2004; Serfling, 2009). Compared to the algorithms based on the LA, the optimization procedure of 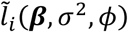, described in a later section, becomes straightforward because the analytical form is available. As zero counts are prevalent in scRNA-seq data, the evaluation of the log-gamma function 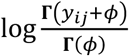 in 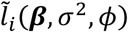 can be saved for most cells. We obtain estimating equations by taking the first derivative of 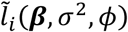 (See Supplementary materials A.4), through which we show in Supplementary materials A.5 that 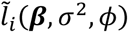 provides an asymptotically consistent estimate of (***β***, *σ*^2^, *ϕ*) when *n*_*i*_ → ∞ and *m* → ∞. The consistency is also supported by the results in our simulation described in the Results section.

As *n*_*i*_ → ∞ is never achieved in real data, we then investigate the performance of using 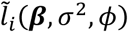 in single-cell data with finite *n*_*i*_. The major question is under which conditions the method works well and how large *n*_*i*_ is required to produce an accurate estimate of *σ*^2^ and *ϕ*. We see from eq. (7) that the accuracy of 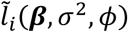 depends on how close 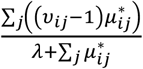 is to zero, and it becomes an exact maximum likelihood estimate (MLE) if this term equals zero. By the Lindeberg central limit theorem (CLT), under mild conditions and the null hypothesis of the fixed effects, 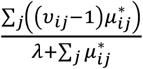 follows a zero-mean normal distribution with a variance at the rate of

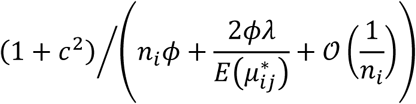

(See Supplementary materials A.6), where *c* is the CV of the scaling factor *π*_*ij*_. Intuitively, the estimation accuracy of NEBULA-LN should depend on the dominant term (1 + *c*^2^)/*n*_*i*_*ϕ* when *n*_*i*_ is large, and this relationship was supported by the simulation study presented in the Results section by comparing the estimates between NEBULA-LN and NEBULA-HL which uses an h-likelihood method (described in the next section). Interestingly, we find that a good estimate of *ϕ* in terms of MSE can be achieved even if 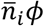 is as small as 20, where 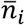 is the CPS value. However, the estimate of *σ*^2^ requires a larger 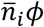 and 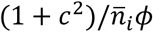 determines the resolution of detecting small *σ*^2^. The simulation study in the Results section shows that 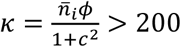 can empirically provide a good estimate of *σ*^2^ as small as ∼0.02. As shown in the analysis of the droplet-based scRNA-seq data (Mathys et al., 2019), only ∼1% of genes, all of which are very low-expressed genes, have very large cell-level overdispersion (*ϕ*^−1^ > 10) in the exitatory neurons. Consequently, NEBULA-LN alone is sufficient for most genes when CPS>200. The standard error (SE) of (***β***, *σ*^2^, *ϕ*) can be obtained by the sandwich covariance estimator estimator (Huber, 1967). We find that the SE based on the Hessian matrix of 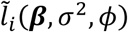 is practically accurate when 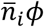 is large. Instead, we choose to use the h-likelihood (described in the next section) for computing its SE.

### Estimation using h-likelihood in NEBULA-HL

As aforementioned, NEBULA-LN is first used to estimate *σ*^2^ and *ϕ*, and if 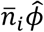 or 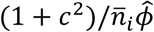 is smaller than a predefined threshold, we then use NEBULA-HL to obtain a more accurate estimate. NEBULA-HL is based on an h-likelihood method (Lee et al., 2006) requiring a two-layer iterative procedure. To derive the h-likelihood, we switch the integral order in *L*_*i*_(***β***, *σ*^2^, *ϕ*) and first integrate out *v*_*ij*_, which leads to

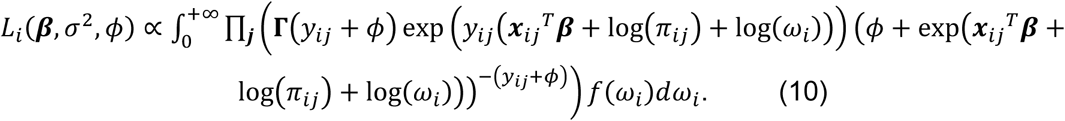

The h-likelihood method first optimizes the integrand in eq. (9) with respect to (***β***, *ω*_*i*_) in the inner loop. Note that *ω*_*i*_ cannot be optimized with ***β*** directly because they are not in the canonical scale (Lee et al., 2006). We thus convert the integrand in eq. (10) to an h-likelihood by re-parametrizing the integral in eq. (9) using *η*_*i*_ = log(*ω*_*i*_), which leads to the log-h-likelihood for individual *i*

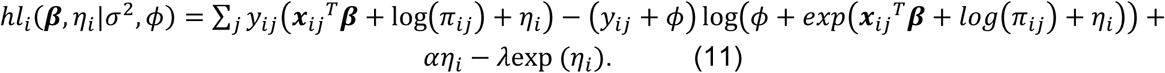

We optimize ∑_*i*_ *hl*_*i*_(***β***, *η*_*i*_|*σ*^2^, *ϕ*) by calculating its first and second derivatives of eq. (11) (See Supplementary materials A.7 for more detail). Because the log-h-likelihood is concave (Both the negative binomial term and the penalizing term are concave), we find that an NR algorithm converges fast and properly. The bottom-right corner of the Hessian matrix corresponding to *η*_*i*_ is diagonal, so the computation for solving the inverse Hessian matrix takes 𝒪(*m*), which is ignorable. We also use this step to compute the SE of ***β*** in NEBULA-LN by plugging in 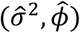 estimated from 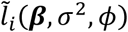.

Given the current estimate (***β***^∗^, *η*_*i*_^∗^), we then optimize (*σ*^2^, *ϕ*) using the marginal likelihood

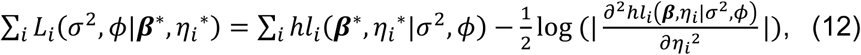

where 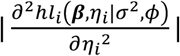 is the logarithm of the absolute value of the determinant of the second derivative with respect to *η*_*i*_. Replacing 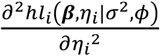 with 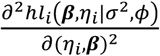 in eq. (12) gives a REML estimate of (*σ*^2^, *ϕ*). Similar to *lme4* (Bates et al., 2014) and *coxmeg* (He and Kulminski, 2020), we use the derivative-free algorithm bobyqa (Powell, 2009) implemented in the *nloptr* R package (Ypma, 2014) for the optimization of ∑_*i*_ *L*_*i*_(*σ*^2^, *ϕ*|***β***^∗^, *η*_*i*_^∗^). The number of iterations in this step grows rapidly with the dimension of the parameters. To reduce the iterations, we plug in 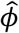 in ∑_*i*_ *L*_*i*_(*σ*^2^, *ϕ*|***β***^∗^, *η*_*i*_^∗^) as we find that 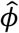 from NEBULA-LN is very accurate under mild conditions (e.g., 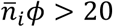). Thus, for most genes estimated by NEBULA-HL, we only perform the optimization for *σ*^2^, which substantially reduces the number of iterations. In most cases, the algorithm can reach convergence within ∼10 iterations.

### Computational implementation of NEBULA-LN

We assessed several algorithms for optimizing 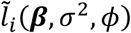 in eq. (8), including (i) the L-BFGS algorithm (Byrd et al., 1995) with box constraints implemented in *nloptr* (Ypma, 2014), (ii) the NR algorithm implemented in the *nlm* R function (Dennis and Schnabel, 1996) and (iii) a trust region algorithm (Fletcher, 1987) implemented in the *trust* R package (https://cran.r-project.org/web/packages/trust/index.html). The NR and the trust region algorithms require the Hessian matrix of 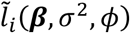. Inspired by (Gilmour et al., 1995), we derive an approximate Hessian matrix (See Supplementary materials A.8) by taking the expectation with respect to *y*_*ij*_ for some elements to simplify the computation. As *σ*^2^, *ϕ* are restricted to be positive values, we reparameterize say what, I guess Hessian matrix using logarithm for the unconstrained optimization. In contrast, the L-BFGS algorithm requires only the estimating equations. We find that the NR algorithm using *nlm* is generally the fastest, and often converges within 10 iterations. However, the NR algorithm may not converge well for low-expressed genes with a small CPC value, and occasionally the convergence depended on the initial values. In contrast, the L-BFGS and trust region algorithms are often slower, but much more robust, and both algorithms produce highly consistent estimates. The trust region algorithm takes fewer iterations, but can be slower than the L-BFGS algorithm when the dimension of ***β*** becomes larger. This is because the computation of the Hessian matrix is more computationally expensive with the increasing number of the parameters. As most genes in single-cell data have low expression or zero counts, the evaluation of most log-gamma, and digamma functions in the marginal likelihood and the estimating equations can be saved or replaced using a recursive formula.

### Comparison with existing tools

We compared the computational performance of NEBULA with three commonly used R packages (*lme4* (Bates et al., 2014), *glmmTMB* (Brooks et al., 2017), and *INLA* (Rue et al., 2009)) for estimating the NBMM. We downloaded the *lme4* and *glmmTMB* R packages via https://cran.r-project.org/ and installed the *INLA* R package via http://www.r-inla.org/download. In *lme4*, we included two methods by setting the argument nAGQ=1 (for LA), and nAGQ=0 (for PQL). In *glmmTMB*, we set family=nbinom2. In *INLA*, we set family = “nbinomial”, and control.predictor = list(compute = FALSE). We kept the other arguments as default. All benchmarks were run in R 3.5 on Windows 10 and Linux, separately.

### Preprocessing of snRNA-seq data

The snRNA-seq UMI count matrix for 17,926 genes and 70,634 cells after QC filtering in 48 human frontal cortex from the ROSMAP project was downloaded from Synapse (https://www.synapse.org/#!Synapse:syn21261143). The details of filtering the cells in the QC step are described in (Mathys et al., 2019). We adopted the clustering results provided by (Mathys et al., 2019), which included 34,976 excitatory neurons, 9196 inhibitory neurons, 18,235 oligodendrocytes, 3392 astrocytes, 1920 microglia, and 2627 OPCs. We did not include pericytes and endothelial cells due to their small sample sizes. We performed a further QC step to remove outlier cells whose total counts or total features were lower than 3 median absolute deviations away from the median using scater (McCarthy et al., 2017), which resulted in 69,458 cells. We also removed genes with CPC<0.1% in the analysis of each cell type (CPC for a gene was calculated by its total UMI counts divided by the number of cells in the cell type). The sub-cell-type clustering annotation provided in (Mathys et al., 2019) was used in the analysis of marker genes for sub-clusters. There were 11, 12, 5, 4, 4, and 3 subclusters in the excitatory neurons, inhibitory neurons, oligodendrocytes, astrocytes, microglia, and OPCs, respectively. The percentage of ribosomal genes (RPS and RPL genes) was computed using the *PercentageFeatureSet* function in the *Seurat* R package (Butler et al., 2018). Subject-level covariates, including age, sex, AD status, race, and *APOE* genotype in the ROSMAP project (Bennett et al., 2012a, 2012b), were downloaded from Synapse (https://www.synapse.org/#!Synapse:syn3157322).

In the co-expression analysis of *APOE*, we first obtained normalized *APOE* expression by dividing the raw count of *APOE* in each cell by its library size. We then subtracted from it the mean value of the normalized *APOE* expression across all cells of each individual. In the isoform-specific analysis, we separated all cells into three categories (e2e3, e3e3, or e4^+^) based on the *APOE* genotypes. We then applied the same normalizing and centering procedure as in the co-expression analysis of *APOE*.

### Simulation study

We generated the number of CPS *n*_*i*_ using a Poisson distribution for the balanced design or a negative binomial distribution with size=3 for the unbalanced design. We evaluated scenarios with the number of subjects *m* = 30, 50, 100 and CPS value 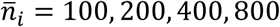. The count data was generated based on the following generative model

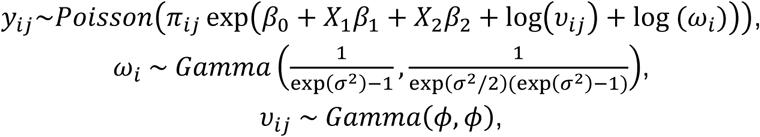

where we considered *β*_0_ ranging from −5 to 2, *ϕ* ranging from 0.01 to 100, and *σ*^2^ ranging from 0.01 to 1. Under the scenarios of a constant scaling factor, we set *π*_*ij*_ = 1, and otherwise we sampled *π*_*ij*_ from *Gamma*(1,1). We simulated a cell-level variable *X*_1_ and a subject-level variable *X*_2_ using a standard normal distribution. Under each of the settings, we generated 500 data sets to calculate the summary statistics. The MSE of an estimate (e.g., 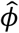) was computed by 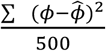 across the 500 data sets.

### Data and code availability

This manuscript was prepared using limited access datasets obtained through Synapse (https://www.synapse.org/#!Synapse:syn3219045, https://www.synapse.org/#!Synapse:syn18485175). The code of this R package can be downloaded and installed via https://github.com/lhe17/nebula.

## Acknowledgments

This research was supported by Grants from the National Institute on Aging (R01 AG061853, R01 AG065477, and R01 AG070488). The funders had no role in study design, data collection and analysis, decision to publish, or manuscript preparation. The content is solely the responsibility of the authors and does not necessarily represent the official views of the National Institutes of Health.

## Author contributions

LH conceived, developed, and implemented the algorithms. LH designed the simulation study and analyzed the simulated and real data. AK contributed to acquiring the real data and discussion of the final results. LH and AK contributed to the writing of the manuscript.

## Supplementary Text

Please see the supplementary file.

## Supplementary Tables

Table S1. A list of top 200 genes co-expressed with the expression of the *APOE* variants at the single-cell level in astrocytes in the human frontal cortex. logFC: The log(fold change) of the gene expression if the *APOE* expression is increased. logFC>0 means a positive correlation, and logFC<0 means a negative correlation with the expression of *APOE*. APOE2: *APOE* e2e3. APOE3: *APOE* e3e3. APOE4: *APOE* e3e4 and *APOE* e4e4.

Table S2. A list of top 200 genes co-expressed with the expression of the *APOE* variants at the single-cell level in microglia in the human frontal cortex. logFC: The log(fold change) of the gene expression if the *APOE* expression is increased. logFC>0 means a positive correlation, and logFC<0 means a negative correlation with the expression of *APOE*. APOE2: *APOE* e2e3. APOE3: *APOE* e3e3. APOE4: *APOE* e3e4 and *APOE* e4e4.

Table S3. The proposed strategy for testing a cell-level predictor used in the analysis of the real data. This table summarizes the condition under which an estimation algorithm (NEBULA-LN or NEBULA-HL) is used to estimate each of overdispersion parameters.

## Supplementary Figures

Figure S1. The computational time (measured in log10(seconds)) of fitting an NBMM for 10,000 genes with respect to the different numbers of fixed-effects predictors in the NBMM. The number of subjects was set at 50, and CPS was set at 100. The average benchmarks are summarized from scenarios of varying subject-level and cell-level overdispersions and the mean count per cell ranging from exp(−4) to 1.

Figure S2. Comparison of estimated cell-level and subject-level overdispersions between NEBULA-LN and NEBULA-HL under a situation in which the CV of the scaling factor (*π*_*ij*_) is one. Five hundred simulated data sets were generated in each of the scenarios to calculate the summary statistics. (A) The cell-level overdispersions estimated by NEBULA-LN and NEBULA-HL under different combinations of CPS, *β*_0_ (a lower *β*_0_ corresponding to a lower CPC value), and *ϕ*. The number of subjects was set at 50. (B) The subject-level overdispersions estimated by NEBULA-LN and NEBULA-HL under different combinations of *β*_0_, *σ*^2^, and *ϕ*. The number of subjects was set at 50. The CPS value was set at 400. (C) The subject-level overdispersions estimated by NEBULA-LN and NEBULA-HL under different combinations of the number of subjects, *σ*^2^, and *ϕ*. The CPS value was set at 400, and *β*_0_ was set at 0.05. (D) The subject-level overdispersions estimated by NEBULA-LN and NEBULA-HL under different combinations of CPS, *σ*^2^, and *ϕ*. The number of subjects was set at 50, and *β*_0_ was set at 1.

Figure S3. Comparison of estimated cell-level overdispersions between NEBULA-LN and NEBULA-HL under different combinations of the number of subjects, *β*_0_, and *ϕ*. The CPS value was set at 400. The summary statistics were calculated from five hundred simulated replicates in each of the scenarios.

Figure S4. The log(FC) of a subject-level predictor estimated by NEBULA versus those estimated by *glmer*.*nb* with nAGQ=0 for 16,207 genes with CPC>0.1% in the excitatory neurons.

Figure S5. Comparison of estimated cell-level and subject-level overdispersions between NEBULA-LN and NEBULA-HL under an unbalanced design in which the cell counts across the subjects were sampled from a negative binomial distribution with size=3. The summary statistics were calculated from five hundred simulated replicates in each of the scenarios. (A) The cell-level overdispersions estimated by NEBULA-LN and NEBULA-HL under different combinations of CPS, *β*_0_ (a lower *β*_0_ corresponding to a lower CPC value), and *ϕ*. The number of subjects was set at 50. (B) The subject-level overdispersions estimated by NEBULA-LN and NEBULA-HL under different combinations of *β*_0_, *σ*^2^, and *ϕ*. The number of subjects was set at 50. The CPS value was set at 400. (C) The subject-level overdispersions estimated by NEBULA-LN and NEBULA-HL under different combinations of the number of subjects, *σ*^2^, and *ϕ*. The CPS value was set at 400, and *β*_0_ was set at 0.05. (D) The subject-level overdispersions estimated by NEBULA-LN and NEBULA-HL under different combinations of CPS, *σ*^2^, and *ϕ*. The number of subjects was set at 50, and *β*_0_ was set at 1.

Figure S6. Empirical type I error rate of testing a subject-level predictor using NEBULA-LN under different combinations of CPC, *σ*^2^, and *ϕ*. The CPS value was set at 400 and the number of subjects was set at 50. The empirical type I error rate was calculated from five hundred simulated replicates in each of the scenarios and was evaluated at the significance level of 0.05 (the dashed lines).

Figure S7. Empirical type I error rate of testing a cell-level predictor using NEBULA-LN under different combinations of CPC, *σ*^2^, and *ϕ*. The number of subjects was set at 50 and the CPS value was set at 400. The empirical type I error rate was calculated from five hundred simulated replicates in each of the scenarios and was evaluated at the significance level of 0.05 (the dashed lines).

Figure S8. Distribution of the library size (the total number of reads) of each cell grouped by the seven major cell types in the snRNA-seq data in the human frontal cortex.

Figure S9. The subject-level overdispersions estimated by NEBULA versus those estimated by *glmer*.*nb* with nAGQ=0 for 16,207 genes with CPC>0.1% in the excitatory neurons.

Figure S10. The cell frequency of the four subclusters in astrocytes across the 48 individuals. The 3386 astrocytes are from the ROSMAP snRNA-seq data in the human frontal cortex.

Figure S11. The cell-level overdispersions of genes with CPC>0.1% in five major cell types in the frontal cortex estimated by NEBULA with respect to the CPC of the gene. No covariates other than the intercept were included in the model.

Figure S12. The subject-level overdispersions of genes with CPC>0.1% in five major cell types in the frontal cortex estimated by NEBULA with respect to the CPC of the gene. No covariates other than the intercept were included in the model.

Figure S13. The distribution of (A) the estimated cell-level overdispersions and (B) subject-level overdispersions in the excitatory neurons (Ex) and the 11 subpopulations in the excitatory neurons (Ex0-Ex12). All genes with CPC>0.1% in each of the cell populations were included.

Figure S14. The cell-level overdispersions estimated by NEBULA versus those estimated by *glmer*.*nb* with nAGQ=0 for 16,207 genes with CPC>0.1% in the excitatory neurons.

Figure S15. Empirical type I error rate of testing a subject-level predictor using PGMM under different combinations of the number of individuals, *σ*^2^, and *ϕ*. The CPS value was set at 200. The empirical type I error rate was calculated from five hundred simulated replicates in each of the scenarios and was evaluated at the significance level of 0.05 (the dashed lines).

