## Supplementary Text for "NEBULA: a fast negative binomial mixed model for differential expression and co-expression analyses of large-scale multi-subject single-cell data"

### A.1. Marginal expectation and variance of the NBGMM

We derive the marginal expectation and variance of $y_{ij}$ under the NBGMM defined in e.q. (5) in the main text. Here, by marginal expectation, we mean the expectation conditional only on $\boldsymbol{x}_{ij}$ and $\pi_{ij}$, but not $\omega_{i}$ or $\upsilon_{ij}$. Based on the model specification, the first and second moments of $y_{ij}$ conditional on $\omega_{i}$ and $\upsilon_{ij}$ are

$$E\left( y_{ij}|\omega_{i},\upsilon_{ij} \right)=\pi_{ij}\exp\left( \boldsymbol{x}_{ij}\boldsymbol{\beta+}\log\left( \omega_{i} \right)+\log\left( \upsilon_{ij} \right) \right)$$

$$E\left( {y_{ij}}^{2}|\omega_{i},\upsilon_{ij} \right)=E\left( y_{ij}|\omega_{i},\upsilon_{ij} \right)+E^{2}\left( y_{ij}|\omega_{i},\upsilon_{ij} \right),$$

where $\boldsymbol{x}_{ij}=(x_{ij1},\ldots,x_{ijk})$ and $\boldsymbol{\beta=}{\boldsymbol{(}\beta_{1}\boldsymbol{,\ldots,}\beta_{k}\boldsymbol{)}}^{T}$. As $\omega_{i}$ follows a gamma distribution defined by e.q. (3), we have its first and second moments

$$E\left( \omega_{i} \right)=\frac{\alpha}{\lambda}=\exp\left( \sigma^{2}/2 \right)$$

$$E\left( {\omega_{i}}^{2} \right)=E^{2}\left( \omega_{i} \right)+Var\left( \omega_{i} \right)=\frac{\alpha^{2}+\alpha}{\lambda^{2}}=\exp\left( 2\sigma^{2} \right).$$

Thus, after taking the expectation over $\upsilon_{ij}$ and $\omega_{i}$ and substituting $E\left( \omega_{i} \right)$, it follows that

$$E\left( y_{ij} \right)=E_{\omega_{i}}\left( E_{\upsilon_{ij}}\left( E\left( y_{ij}|\omega_{i},\upsilon_{ij} \right) \right) \right)$$

$$=E_{\omega_{i}}\left( E_{\upsilon_{ij}}(\upsilon_{ij})\pi_{ij}exp(\boldsymbol{x}_{ij}\boldsymbol{\beta+}\log\left( \omega_{i} \right)) \right)$$

$$=E_{\omega_{i}}\left( \omega_{i}\pi_{ij}exp(\boldsymbol{x}_{ij}\boldsymbol{\beta}) \right)=\pi_{ij}\exp\left( \boldsymbol{x}_{ij}\boldsymbol{\beta+}\frac{\sigma^{2}}{2} \right),$$

and

$$E\left( {y_{ij}}^{2} \right)=E_{\omega_{i}}\left( E_{\upsilon_{ij}}\left( E\left( {y_{ij}}^{2}|\omega_{i},\upsilon_{ij} \right) \right) \right)$$

$$=E\left( y_{ij} \right)+E_{\omega_{i}}\left( E_{\upsilon_{ij}}\left( E^{2}\left( y_{ij}|\omega_{i},\upsilon_{ij} \right) \right) \right)$$

$$=E\left( y_{ij} \right)+E_{\omega_{i}}\left( E_{\upsilon_{ij}}\left( {\upsilon_{ij}}^{2} \right){\pi_{ij}}^{2}\exp\left( 2\left( \boldsymbol{x}_{ij}\boldsymbol{\beta+}\log\left( \omega_{i} \right) \right) \right) \right)$$

$$=E\left( y_{ij} \right)+\left( 1+\frac{1}{\phi} \right)E_{\omega_{i}}\left( {\omega_{i}}^{2}{\pi_{ij}}^{2}\exp\left( 2\boldsymbol{x}_{ij}\boldsymbol{\beta} \right) \right)$$

$$=E\left( y_{ij} \right)+\left( 1+\frac{1}{\phi} \right)\exp\left( 2\sigma^{2} \right){\pi_{ij}}^{2}\exp\left( 2\boldsymbol{x}_{ij}\boldsymbol{\beta} \right)$$

$$=E\left( y_{ij} \right)+\left( 1+\frac{1}{\phi} \right)\exp\left( \sigma^{2} \right)E^{2}\left( y_{ij} \right)$$

Therefore, the marginal variance of $y_{ij}$ is

$Var\left( y_{ij} \right)=E\left( {y_{ij}}^{2} \right)-E^{2}\left( y_{ij} \right)=E\left( y_{ij} \right)+\left( \exp\left( \sigma^{2} \right)-1 \right)E^{2}\left( y_{ij} \right)+\frac{1}{\phi}\exp\left( \sigma^{2} \right)E^{2}\left( y_{ij} \right)$.

### A.2. Proof of convergence in probability

Denote by $\underset{n_{i}\to\infty}{\mathrm{plim}}$ convergence in probability of a sequence to a random variable when $n_{i}\to\infty$. Here, we show that under the condition that $\mu_{ij}^{*}=\exp\left( \boldsymbol{x}_{ij}\boldsymbol{\beta+}\log\left( \pi_{ij} \right) \right)$ has a finite non-zero first moment, we have

$$\underset{n_{i}\to\infty}{\mathrm{plim}}\left( X_{n_{i}}=\frac{\sum_{j=1}^{n_{i}} \left( \upsilon_{ij}-1 \right)\mu_{ij}^{*}}{\lambda+\sum_{j} \mu_{ij}^{*}} \right)=0,$$

that is, given any $\varepsilon>0$, we have$\lim_{n_{i}\to\infty}\Pr\left( {|X}_{n_{i}}|>\varepsilon\right)=0$. Rewrite the sequence $X_{n_{i}}$ as

$X_{n_{i}}=\frac{\frac{\sum_{j=1}^{n_{i}} \left( \upsilon_{ij}-1 \right)\mu_{ij}^{*}}{n_{i}}}{\frac{\lambda+\sum_{j} \mu_{ij}^{*}}{n_{i}}}$.

In fact, as $\upsilon_{ij}$ are *i.i.d.* gamma random variables defined by e.q. (5), and $\mu_{ij}^{*}$ is assumed to be *i.i.d* with a finite first moment, by the weak law of large numbers (WLLN), it follows that

$$\underset{n_{i}\to\infty}{\mathrm{plim}}\frac{\sum_{j=1}^{n_{i}} \left( \upsilon_{ij}-1 \right)\mu_{ij}^{*}}{n_{i}}=E\left( \left( \upsilon_{ij}-1 \right)\mu_{ij}^{*} \right)=E\left( \mu_{ij}^{*} \right)E\left( \upsilon_{ij} \right)-E\left( \mu_{ij}^{*} \right)=0,$$

in which we also use the assumption that the random effects $\upsilon_{ij}$ and the fixed effects $\mu_{ij}^{*}$ are independent. Applying the WLLN again to the denominator gives

$\underset{n_{i}\to\infty}{\mathrm{plim}}\frac{\lambda+\sum_{j} \mu_{ij}^{*}}{n_{i}}=E\left( \mu_{ij}^{*} \right)$,

in which we use $\underset{n_{i}\to\infty}{\mathrm{plim}}\frac{\lambda}{n_{i}}=0$. Thus, it follows by Slutsky's theorem and the assumption $E\left( \mu_{ij}^{*} \right)>0$ that

$$\underset{n_{i}\to\infty}{\mathrm{plim}}\left( X_{n_{i}} \right)=0.$$

### A.3. Approximation of the marginal likelihood using the WLLN

Here we show that the integral for subject $i$ in e.q. (6)

$$I_{i}=\int_{0}^{+\infty} \left( \prod_{j} {\upsilon_{ij}}^{y_{ij}} \right)\underset{(\Theta)}{\underbrace{\left( \lambda+\sum_{j} \left( \upsilon_{ij}\exp\left( \boldsymbol{x}_{ij}\boldsymbol{\beta+}\log\left( \pi_{ij} \right) \right) \right) \right)^{-\left( \sum_{j} y_{ij}+\alpha\right)}}}\prod_{j} f\left( \upsilon_{ij} \right)d\boldsymbol{\upsilon}_{i}$$

can be approximated by e.q. (8)

$$I_{i}\approx\left( \lambda+\sum_{j} \mu_{ij}^{*} \right)^{-\left( \sum_{j} y_{ij}+\alpha\right)}\times\prod_{j=1}^{n_{i}} \int_{0}^{+\infty} {\upsilon_{ij}}^{y_{ij}}\exp\left( -\frac{\left( \sum_{j} y_{ij}+\alpha\right)\left( \upsilon_{ij}-1 \right)\mu_{ij}^{*}}{\lambda+\sum_{j} \mu_{ij}^{*}} \right)\frac{\phi^{\phi}}{\boldsymbol{\Gamma}\left( \phi\right)}{\upsilon_{ij}}^{\phi-1}\exp\left( -\phi\upsilon_{ij} \right)d\upsilon_{ij}$$

under the condition $n_{i}\to\infty$. Rewrite $\Theta$ in e.q.(6) as

$\Theta=\exp\left( -\left( \sum_{j} y_{ij}+\alpha\right)\log\left( \left( \lambda+\sum_{j} \mu_{ij}^{*} \right)\left( 1+\frac{\sum_{j} \left( \left( \upsilon_{ij}-1 \right)\mu_{ij}^{*} \right)}{\lambda+\sum_{j} \mu_{ij}^{*}} \right) \right) \right)$.

As proved in A.2 that $\underset{n_{i}\to\infty}{\mathrm{plim}}\left( \frac{\sum_{j} \left( \left( \upsilon_{ij}-1 \right)\mu_{ij}^{*} \right)}{\lambda+\sum_{j} \mu_{ij}^{*}} \right)=0$, we ignore the contribution to the integral from those $\boldsymbol{\upsilon}_{i}\boldsymbol{\notin}\mathcal{S}$ where $\mathcal{S}:=\left( \boldsymbol{\upsilon}_{i} | \left| \frac{\sum_{j} \left( \left( \upsilon_{ij}-1 \right)\mu_{ij}^{*} \right)}{\lambda+\sum_{j} \mu_{ij}^{*}} \right|<\varepsilon\ll1 \right)$ for a small positive $\varepsilon$ because $P(\boldsymbol{\upsilon}_{i}\boldsymbol{\notin}\mathcal{S)}$ is very small when $n_{i}$ is large. Thus, plugging $f\left( \upsilon_{ij} \right)$ with the gamma density function defined in e.q. (5), we can end up with e.q. (8) as follows

$\int_{0}^{+\infty} \left( \prod_{j} {\upsilon_{ij}}^{y_{ij}} \right)\exp\left( -\left( \sum_{j} y_{ij}+\alpha\right)\log\left( \left( \lambda+\sum_{j} \mu_{ij}^{*} \right)\left( 1+\frac{\sum_{j} \left( \left( \upsilon_{ij}-1 \right)\mu_{ij}^{*} \right)}{\lambda+\sum_{j} \mu_{ij}^{*}} \right) \right) \right)\times\prod_{j=1}^{n_{i}} \frac{\phi^{\phi}}{\boldsymbol{\Gamma}\left( \phi\right)}{\upsilon_{ij}}^{\phi-1}\exp\left( -\phi\upsilon_{ij} \right)d\boldsymbol{\upsilon}_{i}$

$\approx\int_{\boldsymbol{\upsilon}_{i}\mathcal{\in S}} \left( \prod_{j} {\upsilon_{ij}}^{y_{ij}} \right)\exp\left( -\left( \sum_{j} y_{ij}+\alpha\right)\left( \log\left( \lambda+\sum_{j} \mu_{ij}^{*} \right)+\log\left( 1+\frac{\sum_{j} \left( \left( \upsilon_{ij}-1 \right)\mu_{ij}^{*} \right)}{\lambda+\sum_{j} \mu_{ij}^{*}} \right) \right) \right)\times\prod_{j=1}^{n_{i}} \frac{\phi^{\phi}}{\boldsymbol{\Gamma}\left( \phi\right)}{\upsilon_{ij}}^{\phi-1}\exp\left( -\phi\upsilon_{ij} \right)d\boldsymbol{\upsilon}_{i}$

$\approx\int_{\boldsymbol{\upsilon}_{i}\mathcal{\in S}} \left( \prod_{j} {\upsilon_{ij}}^{y_{ij}} \right)\exp\left( -\left( \sum_{j} y_{ij}+\alpha\right)\left( \log\left( \lambda+\sum_{j} \mu_{ij}^{*} \right)+\frac{\sum_{j} \left( \left( \upsilon_{ij}-1 \right)\mu_{ij}^{*} \right)}{\lambda+\sum_{j} \mu_{ij}^{*}} \right) \right)\times\prod_{j=1}^{n_{i}} \frac{\phi^{\phi}}{\boldsymbol{\Gamma}\left( \phi\right)}{\upsilon_{ij}}^{\phi-1}\exp\left( -\phi\upsilon_{ij} \right)d\boldsymbol{\upsilon}_{i}$

$\approx\exp\left( -\left( \sum_{j} y_{ij}+\alpha\right)\log\left( \lambda+\sum_{j} \mu_{ij}^{*} \right) \right)\int_{0}^{+\infty} \left( \prod_{j} {\upsilon_{ij}}^{y_{ij}} \right)\exp\left( -\left( \sum_{j} y_{ij}+\alpha\right)\frac{\sum_{j} \left( \left( \upsilon_{ij}-1 \right)\mu_{ij}^{*} \right)}{\lambda+\sum_{j} \mu_{ij}^{*}} \right)\times\prod_{j=1}^{n_{i}} \frac{\phi^{\phi}}{\boldsymbol{\Gamma}\left( \phi\right)}{\upsilon_{ij}}^{\phi-1}\exp\left( -\phi\upsilon_{ij} \right)d\boldsymbol{\upsilon}_{i}$

$=\left( \lambda+\sum_{j} \mu_{ij}^{*} \right)^{-\left( \sum_{j} y_{ij}+\alpha\right)}\prod_{j=1}^{n_{i}} \int_{0}^{+\infty} {\upsilon_{ij}}^{y_{ij}}\exp\left( \frac{-\left( \sum_{j} y_{ij}+\alpha\right)}{\lambda+\sum_{j} \mu_{ij}^{*}}\left( \upsilon_{ij}-1 \right)\mu_{ij}^{*} \right)\frac{\phi^{\phi}}{\boldsymbol{\Gamma}\left( \phi\right)}{\upsilon_{ij}}^{\phi-1}\exp\left( -\phi\upsilon_{ij} \right)d\upsilon_{ij}$

where the approximations in the second and fourth lines are due to ignoring and adding those Monte Carlo samples $\boldsymbol{\upsilon}_{i}\mathcal{\notin S}$, respectively. The approximation in the third line uses the first-order Taylor expansion of a logarithm function, i.e., $\log\left( 1+x \right)\approx x$ for $x$ close to zero. In the last equation, we change the order between the integral and the product because $\upsilon_{ij}$ are now completely factorized in the integrand. After moving the terms $\exp\left( \frac{\left( \sum_{j} y_{ij}+\alpha\right)\mu_{ij}^{*}}{\lambda+\sum_{j} \mu_{ij}^{*}} \right)$ out of the integral, the integrand is now recognized as a kernel of a gamma density function with respect to $\upsilon_{ij}$. Calculating this integral explicitly gives e.q. (9).

### A.4. The estimating equations in NEBULA-LN

The estimating equations in NEBULA-LN are obtained by taking the first derivative of the approximated marginal likelihood $\sum_{i} \tilde{l}_{i}\left( \boldsymbol{\beta,}\sigma^{2},\phi\right)$ with respect to the parameters $\left( \boldsymbol{\beta,}\sigma^{2},\phi\right)$, where

$\tilde{l}_{i}\left( \boldsymbol{\beta,}\sigma^{2},\phi\right)=\alpha\log\lambda+\log\frac{\boldsymbol{\Gamma}\left( \sum_{j} y_{ij}+\alpha\right)}{\boldsymbol{\Gamma}\left( \alpha\right)}+\sum_{j=1}^{n_{i}} y_{ij}\log\left( \mu_{ij}^{*} \right)-\left( \sum_{j} y_{ij}+\alpha\right)\log\left( \lambda+\sum_{j} \mu_{ij}^{*} \right)+\sum_{j} \left( \phi\log\phi+\log\frac{\boldsymbol{\Gamma}\left( y_{ij}+\phi\right)}{\boldsymbol{\Gamma}\left( \phi\right)}-\left( y_{ij}+\phi\right)\log\left( \phi+\frac{\left( \sum_{j} y_{ij}+\alpha\right)\mu_{ij}^{*}}{\lambda+\sum_{j} \mu_{ij}^{*}} \right)+\frac{\left( \sum_{j} y_{ij}+\alpha\right)\mu_{ij}^{*}}{\lambda+\sum_{j} \mu_{ij}^{*}} \right)$.

To simplify the notations, we define

$$\check{\mu}_{i}:=\lambda+\sum_{j} \mu_{ij}^{*}$$

$$\check{\omega}_{i}:=\frac{\sum_{j} y_{ij}+\alpha}{\check{\mu}_{i}}$$

$\check{\upsilon}_{ij}:=\frac{\phi+y_{ij}}{\phi+\check{\omega}_{i}\mu_{ij}^{*}}$.

We denote by $\alpha^{'}$ and $\lambda^{'}$ the first derivatives of $\alpha$ and $\lambda$ with respect to $\sigma^{2}$, respectively. Through tedious but relatively straightforward calculation, the first derivatives are

$$D_{n_{i},m}^{\boldsymbol{\beta}}=\sum_{i} \frac{\partial\tilde{l}_{i}\left( \boldsymbol{\beta,}\sigma^{2},\phi\right)}{\partial\boldsymbol{\beta}}=\sum_{i} {\boldsymbol{X}_{i}}^{T}\boldsymbol{y}_{i}-\frac{\check{\omega}_{i}}{\check{\mu}_{i}}\left( \sum_{j} \left( 1-\check{\upsilon}_{ij} \right)\mu_{ij}^{*} \right){\boldsymbol{X}_{i}}^{T}\boldsymbol{\mu}_{i}^{*}-\check{\omega}_{i}{\boldsymbol{X}_{i}}^{T}(\boldsymbol{\mu}_{i}^{*}⨀{\check{\boldsymbol{\upsilon}}}_{i})$$

$$D_{n_{i},m}^{\phi}=\sum_{i} \frac{\partial\tilde{l}_{i}\left( \boldsymbol{\beta,}\sigma^{2},\phi\right)}{\partial\phi}=\sum_{i} \sum_{j} \left( \log\phi+\Psi\left( y_{ij}+\phi\right)-\Psi\left( \phi\right)-\log\left( \phi+\check{\omega}_{i}\mu_{ij}^{*} \right)+1-\check{\upsilon}_{ij} \right)$$

$$D_{n_{i},m}^{\sigma^{2}}=\sum_{i} \frac{\partial\tilde{l}_{i}\left( \boldsymbol{\beta,}\sigma^{2},\phi\right)}{\partial\sigma^{2}}=\sum_{i} \alpha^{'}\left( \log\lambda+\Psi\left( \sum_{j} y_{ij}+\alpha\right)-\Psi\left( \alpha\right)-\log\left( \check{\mu}_{i} \right) \right)+\lambda^{'}\left( \frac{\alpha}{\lambda}-\check{\omega}_{i} \right)+\frac{\alpha^{'}-\lambda^{'}\check{\omega}_{i}}{\check{\mu}_{i}}\sum_{j} \left( 1-\check{\upsilon}_{ij} \right)\mu_{ij}^{*}$$

where $\Psi\left( x \right)=\frac{d\log(\boldsymbol{\Gamma}\left( x \right))}{dx}$ is the digamma function, $⨀$ stands for the Hadamard product, $\boldsymbol{y}_{i}={(y_{i1},\ldots,y_{in_{i}})}^{T}$, $\boldsymbol{X}_{i}={({\boldsymbol{x}_{i1}}^{T},\ldots,{\boldsymbol{x}_{in_{i}}}^{T})}^{T}$, $\boldsymbol{\mu}_{i}^{*}={(\mu_{i1}^{*},\ldots,\mu_{in_{i}}^{*})}^{T}$, and ${\check{\boldsymbol{\upsilon}}}_{i}={(\check{\upsilon}_{i1},\ldots,\check{\upsilon}_{in_{i}})}^{T}$. Setting $D_{n_{i},m}^{\boldsymbol{\beta}}$, $D_{n_{i},m}^{\phi}$ and $D_{n_{i},m}^{\sigma^{2}}$ to zero gives a group of estimating equations for $\left( \boldsymbol{\beta,}\sigma^{2},\phi\right)$.

### A.5. Asymptotic consistency of NEBULA-LN

We prove that the estimating equations $D_{n_{i},m}$ given in A.4 lead to a consistent estimator $T_{n_{i},m}$ for $\left( \boldsymbol{\beta,}\sigma^{2},\phi\right)$ when $n_{i}\to\infty$ and $m\to\infty$, that is, we need to show that $T_{n_{i},m}$ asymptotically converges in probability to its true value

$$\underset{\begin{aligned} n_{i}\to\infty\\ m\to\infty\end{aligned}}{\mathrm{plim}}T_{n_{i},m}=\left( \boldsymbol{\beta,}\sigma^{2},\phi\right)$$

The estimator $T_{n_{i},m}$ defined implicitly by the sequence of solutions to the estimation equations $D_{n_{i},m}^{\boldsymbol{\beta,}\sigma^{2},\phi}$ is an M-estimator (Huber, 2004). Asymptotically, the empirical equations $D_{n_{i},m}^{\boldsymbol{\beta,}\sigma^{2},\phi}$ under suitable normalization (divided by $n_{i}$ and $m$) will converge to its deterministic equation $D^{\boldsymbol{\beta,}\sigma^{2},\phi}$ by the WLLN, that is,

$\underset{\begin{aligned} n_{i}\to\infty\\ m\to\infty\end{aligned}}{\mathrm{plim}}D_{n_{i},m}^{\boldsymbol{\beta,}\sigma^{2},\phi}=D^{\boldsymbol{\beta,}\sigma^{2},\phi}$

(see e.g., (Serfling, 2009) section 7.2). Under regularity conditions that $D_{n_{i},m}^{\boldsymbol{\beta,}\sigma^{2},\phi}$ is continuous and dominated by an integrable function within a compact parameter space containing the true values (see e.g., (Serfling, 2009) section 7.2 or (Vaart, 1998) section 5.2), the asymptotical consistency of $T_{n_{i},m}$ amounts to proving that the true values of $\left( \boldsymbol{\beta,}\sigma^{2},\phi\right)$ are solutions to $D^{\boldsymbol{\beta,}\sigma^{2},\phi}=0$.

We begin by examing the term $1-\check{\upsilon}_{ij}$ present in all the estimating equations. By the WLLN and $E\left( y_{ij}|\mu_{ij}^{*},\omega_{i} \right)=\omega_{i}\mu_{ij}^{*}$ in A.1, it follows that

$\underset{n_{i}\to\infty}{\mathrm{plim}}\check{\omega}_{i}=\underset{n_{i}\to\infty}{\mathrm{plim}}\frac{\frac{\sum_{j} y_{ij}+\alpha}{n_{i}}}{\frac{\lambda+\sum_{j} \mu_{ij}^{*}}{n_{i}}}=\frac{\omega_{i}E\left( \mu_{ij}^{*} \right)}{E\left( \mu_{ij}^{*} \right)}=\omega_{i}$.

Then because $\check{\upsilon}_{ij}$ is continuous uniformly in $y_{ij}$ and has a finite first moment, by Lemma 7.2.2A in (Serfling, 2009), we have

$$\underset{n_{i}\to\infty}{\mathrm{plim}}\frac{\sum_{j} \check{\upsilon}_{ij}}{n_{i}}=\frac{\phi+E(y_{ij}|\mu_{ij}^{*},\omega_{i})}{\phi+\omega_{i}\mu_{ij}^{*}}=\frac{\phi+\omega_{i}\mu_{ij}^{*}}{\phi+\omega_{i}\mu_{ij}^{*}}=1.$$

Hence, all terms involving $1-\check{\upsilon}_{ij}$ cancel out.

Following (Sutradhar and Qu, 1998),

$$\underset{m\to\infty}{\mathrm{plim}}\frac{\sum_{i} \check{\omega}_{i}}{m}=E\left( \check{\omega}_{i} \right)=E\left( E\left( \check{\omega}_{i}|\omega_{i} \right) \right)$$

$$=E\left( \frac{\sum_{j} E\left( y_{ij} | \omega_{i} \right)+\alpha}{\lambda+\sum_{j} \mu_{ij}^{*}} \right)=E\left( \frac{\omega_{i}\sum_{j} \mu_{ij}^{*}+\alpha}{\lambda+\sum_{j} \mu_{ij}^{*}} \right)=\frac{\frac{\alpha}{\lambda}\sum_{j} \mu_{ij}^{*}+\alpha}{\lambda+\sum_{j} \mu_{ij}^{*}}=\frac{\alpha}{\lambda}.$$

Hence, the term $\frac{\alpha}{\lambda}-\check{\omega}_{i}$ in $D_{n_{i},m}^{\sigma^{2}}$ cancels out. In $\underset{\begin{aligned} n_{i}\to\infty\\ m\to\infty\end{aligned}}{\mathrm{plim}}D_{n_{i},m}^{\boldsymbol{\beta}}$, it remains to show that under suitable normalization,

$$\underset{n_{i}\to\infty}{\mathrm{plim}}{\boldsymbol{X}_{i}}^{T}\boldsymbol{y}_{i}-\check{\omega}_{i}{\boldsymbol{X}_{i}}^{T}\left( \boldsymbol{\mu}_{i}^{*}⨀{\check{\boldsymbol{\upsilon}}}_{i} \right)=0.$$

In fact, for the *k*th component in $\boldsymbol{\beta}$, we have for the first term

$$\underset{n_{i}\to\infty}{\mathrm{plim}}\frac{{\boldsymbol{X}_{i}}^{T}\boldsymbol{y}_{i}}{n_{i}}=\underset{n_{i}\to\infty}{\mathrm{plim}}\frac{\sum_{j} x_{ijk}y_{ij}}{n_{i}}=x_{ijk}E\left( y_{ij} \right)=\omega_{i}x_{ijk}\mu_{ij}^{*}$$

and for the second term by the continuous mapping theorem, Slutsky's theorem and Lemma 7.2.2A in (Serfling, 2009),

$$\underset{n_{i}\to\infty}{\mathrm{plim}}\frac{{\boldsymbol{X}_{i}}^{T}\check{\omega}_{i}\left( \boldsymbol{\mu}_{i}^{*}⨀{\check{\boldsymbol{\upsilon}}}_{i} \right)}{n_{i}}=\underset{n_{i}\to\infty}{\mathrm{plim}}\check{\omega}_{i}\cdot\underset{n_{i}\to\infty}{\mathrm{plim}}\frac{\sum_{j} x_{ijk}\mu_{ij}^{*}\check{\upsilon}_{ij}}{n_{i}}=\omega_{i}\frac{x_{ijk}\mu_{ij}^{*}\left( \phi+E\left( y_{ij} | \mu_{ij}^{*},\omega_{i} \right) \right)}{\phi+\omega_{i}\mu_{ij}^{*}}=\omega_{i}x_{ijk}\mu_{ij}^{*}$$

which proves that $\underset{\begin{aligned} n_{i}\to\infty\\ m\to\infty\end{aligned}}{\mathrm{plim}}D_{n_{i},m}^{\boldsymbol{\beta}}=0$.

In $D_{n_{i},m}^{\sigma^{2}}$, it remains to show that $\underset{\begin{aligned} n_{i}\to\infty\\ m\to\infty\end{aligned}}{\mathrm{plim}}\log\lambda-\Psi\left( \alpha\right)+\Psi\left( \sum_{j} y_{ij}+\alpha\right)-\log\left( \check{\mu}_{i} \right)=0$. As $\omega_{i}$ follows a gamma distribution defined by e.q. (3), it can be verified based on the density function of $log(\omega_{i})$ (also see (Sutradhar and Qu, 1998)) that its expectation is

$E\left( \log\left( \omega_{i} \right) \right)=\Psi\left( \alpha\right)-\log\lambda$.

Now consider the posterior distribution of $\omega_{i}$ conditional on $\boldsymbol{y}_{i}$ and $\boldsymbol{\upsilon}_{i}$, which can be shown as

$$f\left( \omega_{i} | \boldsymbol{y}_{i},\boldsymbol{\upsilon}_{i} \right)$$

$$\propto f\left( \boldsymbol{y}_{i} | \omega_{i},\boldsymbol{\upsilon}_{i} \right)f\left( \omega_{i} \right)$$

$$=\prod_{j} Pois\left( y_{ij}|\omega_{i}\upsilon_{ij}\mu_{ij}^{*} \right)Gamma\left( \omega_{i}|\alpha,\lambda\right)$$

$=Gamma\left( \alpha+\sum_{j} y_{ij},\lambda+\sum_{j} \upsilon_{ij}\mu_{ij}^{*} \right)$.

Therefore, it follows that

$$E\left( \log\left( \omega_{i} \right) | \boldsymbol{y}_{i},\boldsymbol{\upsilon}_{i} \right)=\Psi\left( \sum_{j} y_{ij}+\alpha\right)-\log\left( \lambda+\sum_{j} \upsilon_{ij}\mu_{ij}^{*} \right)$$

and

$$E\left( \log\left( \omega_{i} \right) | \boldsymbol{y}_{i} \right)=E_{\boldsymbol{\upsilon}_{i}}\left( E\left( \log\left( \omega_{i} \right) | \boldsymbol{y}_{i},\boldsymbol{\upsilon}_{i} \right) \right)=\Psi\left( \sum_{j} y_{ij}+\alpha\right)-E_{\boldsymbol{\upsilon}_{i}}\log\left( \lambda+\sum_{j} \upsilon_{ij}\mu_{ij}^{*} \right)$$

Under the limit $m\to\infty$, by substituting $E\left( \log\left( \omega_{i} \right) | \boldsymbol{y}_{i} \right)$, we have

$$\underset{m\to\infty}{\mathrm{plim}}\frac{\sum_{i} \Psi\left( \sum_{j} y_{ij}+\alpha\right)-\log\check{\mu}_{i}}{m}$$

$$=\underset{m\to\infty}{\mathrm{plim}}\frac{\sum_{i} \Psi\left( \sum_{j} y_{ij}+\alpha\right)-E_{\boldsymbol{\upsilon}_{i}}\log\left( \lambda+\sum_{j} \upsilon_{ij}\mu_{ij}^{*} \right)+E_{\boldsymbol{\upsilon}_{i}}\log\left( \lambda+\sum_{j} \upsilon_{ij}\mu_{ij}^{*} \right)-\log\check{\mu}_{i}}{m}$$

$$=\underset{m\to\infty}{\mathrm{plim}}\frac{\sum_{i} \Psi\left( \sum_{j} y_{ij}+\alpha\right)-E_{\boldsymbol{\upsilon}_{i}}\log\left( \lambda+\sum_{j} \upsilon_{ij}\mu_{ij}^{*} \right)}{m}+\lim_{m\to\infty}\frac{\sum_{i} E_{\boldsymbol{\upsilon}_{i}}\log\left( \lambda+\sum_{j} \upsilon_{ij}\mu_{ij}^{*} \right)-\log\check{\mu}_{i}}{m}$$

$$=E_{\boldsymbol{y}_{i}}\left( E\left( \log\left( \omega_{i} \right) | \boldsymbol{y}_{i} \right) \right)+\lim_{m\to\infty}\frac{\sum_{i} E_{\boldsymbol{\upsilon}_{i}}\log\left( \frac{\lambda+\sum_{j} \upsilon_{ij}\mu_{ij}^{*}}{\check{\mu}_{i}} \right)}{m}=E\left( \log\left( \omega_{i} \right) \right).$$

The second term in the above equation converges to zero under $n_{i}\to\infty$, that is,

$$\lim_{n_{i}\to\infty}E_{\boldsymbol{\upsilon}_{i}}\log\left( \frac{\lambda+\sum_{j} \upsilon_{ij}\mu_{ij}^{*}}{\check{\mu}_{i}} \right)=0.$$

This is because by WLLN, Slutsky's theorem and the continuous mapping theorem, we have

$$\underset{n_{i}\to\infty}{\mathrm{plim}}\log\left( \frac{\lambda+\sum_{j} \upsilon_{ij}\mu_{ij}^{*}}{\check{\mu}_{i}} \right)=\underset{n_{i}\to\infty}{\mathrm{plim}}\log\left( \frac{\lambda+\sum_{j} \upsilon_{ij}\mu_{ij}^{*}}{\lambda+\sum_{j} \mu_{ij}^{*}} \right)=\underset{n_{i}\to\infty}{\mathrm{plim}}\log\left( \frac{\frac{\lambda+\sum_{j} \upsilon_{ij}\mu_{ij}^{*}}{n_{i}}}{\frac{\lambda+\sum_{j} \mu_{ij}^{*}}{n_{i}}} \right)=0$$

and $\frac{\lambda+\sum_{j} \upsilon_{ij}\mu_{ij}^{*}}{\check{\mu}_{i}}$ is uniformly integrable as it has a finite second moment. By Theorem 1.4A in (Serfling, 2009), convergence in probability implies convergence in mean in this case. Hence, we end up with $\underset{\begin{aligned} n_{i}\to\infty\\ m\to\infty\end{aligned}}{\mathrm{plim}}D_{n_{i},m}^{\sigma^{2}}=0$.

Finally, we show that $\underset{\begin{aligned} n_{i}\to\infty\\ m\to\infty\end{aligned}}{\mathrm{plim}}\log\phi+\Psi\left( y_{ij}+\phi\right)-\Psi\left( \phi\right)-\log\left( \phi+\check{\omega}_{i}\mu_{ij}^{*} \right)=0$ in $D_{n_{i},m}^{\phi}$. As $\upsilon_{ij}$ follows a gamma distribution defined by e.q. (3), it can be verified that

$$E\left( \log\left( \upsilon_{ij} \right) \right)=\Psi\left( \phi\right)-\log\phi.$$

Again, consider the conditional posterior distribution

$$f\left( \upsilon_{ij} | y_{ij},\omega_{i} \right)$$

$$\propto f\left( y_{ij} | \omega_{i},\upsilon_{ij} \right)f\left( \upsilon_{ij} \right)$$

$$=Pois\left( y_{ij}|\omega_{i}\upsilon_{ij}\mu_{ij}^{*} \right)Gamma\left( \upsilon_{ij}|\phi,\phi\right)$$

$$=Gamma\left( y_{ij}+\phi,\omega_{i}\mu_{ij}^{*}+\phi\right).$$

Therefore, conditional on $\omega_{i}$, we have

$$E\left( \log\left( \upsilon_{ij} \right)|y_{ij},\omega_{i} \right)=\Psi\left( y_{ij}+\phi\right)-\log\left( \phi+\omega_{i}\mu_{ij}^{*} \right),$$

and

$$E\left( \log\left( \upsilon_{ij} \right)|y_{ij} \right)=E_{\omega_{i}}\left( E\left( \log\left( \upsilon_{ij} \right)|y_{ij},\omega_{i} \right) \right)=\Psi\left( y_{ij}+\phi\right)-E_{\omega_{i}}\left( \log\left( \phi+\omega_{i}\mu_{ij}^{*} \right) \right)$$

Now by the continuous mapping theorem and plugging in the result $\underset{n_{i}\to\infty}{\mathrm{plim}}\check{\omega}_{i}=\omega_{i}$, it follows that

$$\underset{\begin{aligned} n_{i}\to\infty\\ m\to\infty\end{aligned}}{\mathrm{plim}}\frac{1}{m}\sum_{i} \frac{\sum_{j} \Psi\left( y_{ij}+\phi\right)-\log\left( \phi+\check{\omega}_{i}\mu_{ij}^{*} \right)}{n_{i}}$$

$$=\underset{\begin{aligned} n_{i}\to\infty\\ m\to\infty\end{aligned}}{\mathrm{plim}}\frac{1}{m}\sum_{i} \frac{\sum_{j} \Psi\left( y_{ij}+\phi\right)-\log\left( \phi+\omega_{i}\mu_{ij}^{*} \right)}{n_{i}}.$$

Applying the WLLN to the first summation over $\omega_{i}$ and then to the second summation over $y_{ij}$, it follows that

$$\underset{\begin{aligned} n_{i}\to\infty\\ m\to\infty\end{aligned}}{\mathrm{plim}}\frac{1}{m}\sum_{i} \frac{\sum_{j} \Psi\left( y_{ij}+\phi\right)-\log\left( \phi+\omega_{i}\mu_{ij}^{*} \right)}{n_{i}}$$

$$=\underset{n_{i}\to\infty}{\mathrm{plim}}E_{\omega_{i}}\frac{\sum_{j} \Psi\left( y_{ij}+\phi\right)-\log\left( \phi+\omega_{i}\mu_{ij}^{*} \right)}{n_{i}}$$

$$=E_{\omega_{i}}\left( E_{y_{ij}}\left( \Psi\left( y_{ij}+\phi\right)-\log\left( \phi+\omega_{i}\mu_{ij}^{*} \right) \right) \right)$$

$$=E_{\omega_{i}}\left( E_{y_{ij}}\left( E\left( \log\left( \upsilon_{ij} \right)|y_{ij},\omega_{i} \right) \right) \right)=E\left( \log\left( \upsilon_{ij} \right) \right)$$

This completes the proof of the asymptotical consistency of the estimator $T_{n_{i},m}$.

### A.6. Proof of asymptotic normality

We show that under mild conditions and the null model, $X_{n_{i}}=\frac{\sum_{j} \left( \left( \upsilon_{ij}-1 \right)\mu_{ij}^{*} \right)}{\lambda+\sum_{j} \mu_{ij}^{*}}$ asymptotically follows a zero-mean normal distribution with a variance at the rate of $1/\left( n_{i}\phi+\frac{2\phi\lambda}{E\left( \mu_{ij}^{*} \right)}+O\left( \frac{1}{n_{i}} \right) \right)$. In A.1., we have shown that

$$\underset{n_{i}\to\infty}{\mathrm{plim}}\left( X_{n_{i}}=\frac{\sum_{j=1}^{n_{i}} \left( \upsilon_{ij}-1 \right)\mu_{ij}^{*}}{\lambda+\sum_{j} \mu_{ij}^{*}} \right)=0.$$

Hence, it remains to prove the asymptotical normality of $X_{n_{i}}$ and show that the variance of $X_{n_{i}}$ is $1/\left( n_{i}\phi+\frac{2\phi\lambda}{E\left( \mu_{ij}^{*} \right)}+O\left( \frac{1}{n_{i}} \right) \right)$. As $\upsilon_{ij}$ is *i.i.d* gamma random variable, by the Lindeberg central limit theorem, when $n_{i}\to\infty$ and the Lindeberg’s condition is satisfied, we have

$$\frac{\sum_{j} \left( \left( \upsilon_{ij}-1 \right)\mu_{ij}^{*} \right)}{\sqrt{\sum_{j} \left( Var\left( \left( \upsilon_{ij}-1 \right)\mu_{ij}^{*} \right) \right)}}\underset{\to}{d}\mathcal{N}\left( 0,1 \right),$$

where $\underset{\to}{d}$ denotes convergence in distribution. Then, plugging in $Var\left( \upsilon_{ij} \right)=1/\phi$, it follows that

$$X_{n_{i}}=\frac{\sum_{j} \left( \left( \upsilon_{ij}-1 \right)\mu_{ij}^{*} \right)}{\lambda+\sum_{j} \mu_{ij}^{*}}$$

$$=\frac{\sqrt{\sum_{j} \left( Var\left( \left( \upsilon_{ij}-1 \right)\mu_{ij}^{*} \right) \right)}}{\lambda+\sum_{j} \mu_{ij}^{*}}\frac{\sum_{j} \left( \left( \upsilon_{ij}-1 \right)\mu_{ij}^{*} \right)}{\sqrt{\sum_{j} \left( Var\left( \left( \upsilon_{ij}-1 \right)\mu_{ij}^{*} \right) \right)}}$$

$$\underset{\to}{d}N\left( 0,\tau_{X_{n_{i}}}^{2}=\frac{\sum_{j} \left( Var\left( \left( \upsilon_{ij}-1 \right)\mu_{ij}^{*} \right) \right)}{\left( \lambda+\sum_{j} \mu_{ij}^{*} \right)^{2}} \right)$$

$$=N\left( 0,\tau_{X_{n_{i}}}^{2}=\frac{\sum_{j} {\mu_{ij}^{*}}^{2}}{\phi\left( \lambda+\sum_{j} \mu_{ij}^{*} \right)^{2}} \right).$$

where $\tau_{X_{n_{i}}}^{2}$ is the variance of the normal distribution. For simplicity but without loss of generality, we consider the null model where $\mu_{ij}^{*}=\pi_{ij}\exp\left( \beta_{0} \right)$ has only the intercept term and all variation of $y_{ij}$ is included in $\phi$ and $\sigma^{2}$. In this case, $\tau_{X_{n_{i}}}^{2}$ can be expressed as

$$\tau_{X_{n_{i}}}^{2}=\frac{\exp\left( 2\beta_{0} \right)\sum_{j} {\pi_{ij}}^{2}}{\phi\left( \lambda^{2}+2\lambda\exp\left( \beta_{0} \right)\sum_{j} \pi_{ij}+\exp\left( 2\beta_{0} \right){(\sum_{j} \pi_{ij})}^{2} \right)}.$$

Denote by $\pi=E\left( \pi_{ij} \right)$ the mean of the scaling factor, and by $c=\frac{\sqrt{Var\left( \pi_{ij} \right)}}{\pi}$ the coefficient of variation of the scaling factor. We found that $c^{2}$ is often small (~0.25) in most cell types (microglia, oligodendrocytes, astrocytes, and OPCs) in e.g., the snRNA-seq data in (Mathys et al., 2019) when the total library size of a cell is used as the scaling factor. In excitatory and inhibitory neurons, we observed $c^{2}\approx0.8$, probably because different types of neurons varied significantly in terms of morphology. Then, when $n_{i}$ is large, we can approximately rewrite $\tau_{X_{n_{i}}}^{2}$ by plugging in $\pi$ and $c$ as

$$\tau_{X_{n_{i}}}^{2}=\frac{\frac{\exp\left( 2\beta_{0} \right)\sum_{j} {\pi_{ij}}^{2}}{{n_{i}}^{2}}}{\frac{\phi\left( \lambda^{2}+2\lambda\exp\left( \beta_{0} \right)\sum_{j} \pi_{ij}+\exp\left( 2\beta_{0} \right)\left( \sum_{j} \pi_{ij} \right)^{2} \right)}{{n_{i}}^{2}}}$$

$$\approx\frac{\frac{\exp\left( 2\beta_{0} \right)\left( Var\left( \pi_{ij} \right)+\pi^{2} \right)}{n_{i}}}{\phi\left( \frac{\lambda^{2}}{{n_{i}}^{2}}+\frac{2\lambda\exp\left( \beta_{0} \right)\pi}{n_{i}}+\exp\left( 2\beta_{0} \right)\pi^{2} \right)}$$

$$=\frac{1}{\frac{\phi n_{i}}{1+c^{2}}+\frac{2\lambda\phi}{\left( 1+c^{2} \right)\exp\left( \beta_{0} \right)\pi}+\frac{\phi\lambda^{2}}{\left( 1+c^{2} \right)\exp\left( 2\beta_{0} \right)\pi^{2}n_{i}}}$$

$$=\frac{\left( 1+c^{2} \right)}{\phi n_{i}+\frac{2\lambda\phi}{\exp\left( \beta_{0} \right)\pi}+\frac{\phi\lambda^{2}}{\exp\left( 2\beta_{0} \right)\pi^{2}n_{i}}}.$$

Thus, the leading term in $\tau_{X_{n_{i}}}^{2}$is $\frac{\phi n_{i}}{1+c^{2}}$ when $n_{i}\to\infty$. It is also clear that the second term can be large for low-expressed genes because of $\exp\left( \beta_{0} \right)\pi\ll1$.

### A.7. The Newton-Raphson algorithm for optimizing the h-likelihood

Here, we derive the NR algorithm for optimization of the h-likelihood in NEBULA-HL, which requires calculating the first and second derivatives of the h-likelihood. After parametrizing using $\eta_{i}=log \left( \omega_{i} \right)$ in e.q. (10), we obtain the following h-likelihood

$$hl\left( \boldsymbol{\beta,\eta}|\sigma^{2},\phi\right)=\sum_{i} {hl}_{i}\left( \boldsymbol{\beta,}\eta_{i}|\sigma^{2},\phi\right)=\sum_{i} \sum_{j} y_{ij}\left( \boldsymbol{x}_{ij}\boldsymbol{\beta+}\log\left( \pi_{ij} \right)\boldsymbol{+}\eta_{i} \right)-\left( y_{ij}+\phi\right)\log\left( \phi+\exp\left( \boldsymbol{x}_{ij}\boldsymbol{\beta+}\log\left( \pi_{ij} \right)\boldsymbol{+}\eta_{i} \right) \right)+\alpha\eta_{i}-\lambda\exp\left( \eta_{i} \right),$$

in which $\boldsymbol{\beta}$ and $\boldsymbol{\eta}$ are now on the canonical scale, and thus can be optimized simultaneously. Taking the first derivative with respect to $\boldsymbol{\beta}$ and $\eta_{i}$ gives

$$\frac{\partial hl\left( \boldsymbol{\beta,\eta}|\sigma^{2},\phi\right)}{\partial\boldsymbol{\beta}}=\sum_{i} \sum_{j} \left( y_{ij}-\frac{\left( y_{ij}+\phi\right)\exp\left( \boldsymbol{x}_{ij}\boldsymbol{\beta+}\log\left( \pi_{ij} \right)\boldsymbol{+}\eta_{i} \right)}{\phi+\exp\left( \boldsymbol{x}_{ij}\boldsymbol{\beta+}\log\left( \pi_{ij} \right)\boldsymbol{+}\eta_{i} \right)} \right)\boldsymbol{x}_{ij}$$

$$\frac{\partial hl\left( \boldsymbol{\beta,\eta}|\sigma^{2},\phi\right)}{\partial\eta_{i}}=\sum_{j} \left( y_{ij}-\frac{\left( y_{ij}+\phi\right)\exp\left( \boldsymbol{x}_{ij}\boldsymbol{\beta+}\log\left( \pi_{ij} \right)\boldsymbol{+}\eta_{i} \right)}{\phi+\exp\left( \boldsymbol{x}_{ij}\boldsymbol{\beta+}\log\left( \pi_{ij} \right)\boldsymbol{+}\eta_{i} \right)} \right)+\alpha-\lambda\exp\left( \eta_{i} \right).$$

Taking the derivative of $\frac{\partial{hl}_{i}\left( \boldsymbol{\beta,}\eta_{i}|\sigma^{2},\phi\right)}{\partial\boldsymbol{\beta}}$ and $\frac{\partial{hl}_{i}\left( \boldsymbol{\beta,}\eta_{i}|\sigma^{2},\phi\right)}{\partial\eta_{i}}$ again with respect to $\boldsymbol{\beta}$ and $\eta_{i}$ gives the Hessian matrix $\boldsymbol{H}$ with the following entries

$$\frac{\partial^{2}hl\left( \boldsymbol{\beta,\eta}|\sigma^{2},\phi\right)}{\partial\beta_{s}\partial\beta_{t}}=\sum_{i} \sum_{j} -z_{ij}x_{ijs}x_{ijt}$$

$$\frac{\partial^{2}hl\left( \boldsymbol{\beta,\eta}|\sigma^{2},\phi\right)}{\partial{\eta_{i}}^{2}}=-\lambda\exp\left( \eta_{i} \right)-\sum_{j} z_{ij}$$

$$\frac{\partial^{2}hl\left( \boldsymbol{\beta,\eta}|\sigma^{2},\phi\right)}{\partial\beta_{s}\partial\eta_{i}}=\sum_{j} -z_{ij}x_{ijs}\eta_{i}$$

where $z_{ij}=\frac{\phi\left( y_{ij}+\phi\right)\exp\left( \boldsymbol{x}_{ij}\boldsymbol{\beta+}\log\left( \pi_{ij} \right)\boldsymbol{+}\eta_{i} \right)}{\left( \phi+\exp\left( \boldsymbol{x}_{ij}\boldsymbol{\beta+}\log\left( \pi_{ij} \right)\boldsymbol{+}\eta_{i} \right) \right)^{2}}$. And all second derivatives $\frac{\partial^{2}{hl}_{i}\left( \boldsymbol{\beta,}\eta_{i}|\sigma^{2},\phi\right)}{\partial\eta_{i}\partial\eta_{j}}$ equal zero. Therefore, $\boldsymbol{H}$ is highly sparse and the bottom-right block matrix is diagonal. Then, an NR algorithm updates $(\boldsymbol{\beta,\eta)}$ at each iteration using

$${(\boldsymbol{\beta}^{new}\boldsymbol{,}\boldsymbol{\eta}^{new}\boldsymbol{)}}^{T}={(\boldsymbol{\beta}^{old}\boldsymbol{,}\boldsymbol{\eta}^{old}\boldsymbol{)}}^{T}-\boldsymbol{H}^{-1}\left( \frac{\partial hl\left( \boldsymbol{\beta,\eta}|\sigma^{2},\phi\right)}{\partial\boldsymbol{\beta}}\boldsymbol{,}\frac{\partial hl\left( \boldsymbol{\beta,\eta}|\sigma^{2},\phi\right)}{\partial\eta_{i}} \right)^{T}$$

until the increase of the h-likelihood is lower than a pre-defined tolerance and reaches convergence.

### A.8. The Hessian matrix for the Newton-Raphson algorithm in NEBULA-LN

We derive an NR algorithm for optimizing $\tilde{l}_{i}\left( \boldsymbol{\beta,}\sigma^{2},\phi\right)$ in NEBULA-LN. The first derivatives of $\tilde{l}_{i}\left( \boldsymbol{\beta,}\sigma^{2},\phi\right)$ are given in A.4. It remains to calculate the Hessian matrix by taking the second derivatives. We use an approximation for some terms by taking the expectation over $y_{ij}$ to simply the results. We define $\phi_{ij}^{*}:=\phi+\check{\omega}_{i}\mu_{ij}^{*}$, ${\bar{\boldsymbol{x}}}_{i}:={\boldsymbol{\mu}_{i}^{*}}^{T}\boldsymbol{X}_{i}$, and ${\bar{\boldsymbol{X}}}_{i}:={\boldsymbol{X}_{i}}^{T}\mathrm{diag}\boldsymbol{(}\boldsymbol{\mu}_{i}^{*}\boldsymbol{)}\boldsymbol{X}_{i}$. We denote by $\alpha^{''}$ and $\lambda^{''}$ the second derivative of $\alpha$ and $\lambda$ with respect to $\sigma^{2}$. Through some calculation, it follows that

$$\frac{\partial^{2}\tilde{l}_{i}\left( \boldsymbol{\beta,}\sigma^{2},\phi\right)}{\partial\beta_{s}\partial\beta_{t}}=-\frac{{\check{\omega}_{i}}^{2}\bar{x}_{is}}{\check{\mu}_{i}}\sum_{j} \frac{\check{\upsilon}_{ij}{\mu_{ij}^{*}}^{2}x_{ijt}}{\phi_{ij}^{*}}-\frac{{\check{\omega}_{i}}^{2}\bar{x}_{it}}{\check{\mu}_{i}}\sum_{j} \frac{\check{\upsilon}_{ij}{\mu_{ij}^{*}}^{2}x_{ijs}}{\phi_{ij}^{*}}+\frac{{\check{\omega}_{i}}^{2}\bar{x}_{is}\bar{x}_{it}}{{\check{\mu}_{i}}^{2}}\sum_{j} \frac{\check{\upsilon}_{ij}{\mu_{ij}^{*}}^{2}}{\phi_{ij}^{*}}-\check{\omega}_{i}\sum_{j} \check{\upsilon}_{ij}\mu_{ij}^{*}x_{ijs}x_{ijt}\boldsymbol{+}\frac{\check{\omega}_{i}\bar{x}_{it}}{\check{\mu}_{i}}\sum_{j} \check{\upsilon}_{ij}\mu_{ij}^{*}x_{ijs}+\frac{\check{\omega}_{i}\bar{x}_{is}}{\check{\mu}_{i}}\sum_{j} \check{\upsilon}_{ij}\mu_{ij}^{*}x_{ijt}+{\check{\omega}_{i}}^{2}\sum_{j} \frac{\check{\upsilon}_{ij}{\mu_{ij}^{*}}^{2}x_{ijs}x_{ijt}}{\phi_{ij}^{*}}-\frac{\check{\omega}_{i}\bar{x}_{is}\bar{x}_{it}}{\check{\mu}_{i}}+\frac{\check{\omega}_{i}{\bar{\boldsymbol{X}}}_{ist}}{\check{\mu}_{i}}\sum_{j} \left( 1-\check{\upsilon}_{ij} \right)\mu_{ij}^{*}+\frac{2\check{\omega}_{i}\bar{x}_{is}\bar{x}_{it}}{\check{\mu}_{i}}\sum_{j} \left( 1-\check{\upsilon}_{ij} \right)\mu_{ij}^{*}$$

$$\frac{\partial^{2}\tilde{l}_{i}\left( \boldsymbol{\beta,}\sigma^{2},\phi\right)}{\partial\phi^{2}}=\sum_{j} \frac{1}{\phi}+\Psi^{'}\left( y_{ij}+\phi\right)-\Psi^{'}\left( \phi\right)-\frac{2-\check{\upsilon}_{ij}}{\phi_{ij}^{*}}$$

$$\frac{\partial^{2}\tilde{l}_{i}\left( \boldsymbol{\beta,}\sigma^{2},\phi\right)}{\partial\sigma^{4}}=\alpha^{''}\log\lambda+\frac{2\alpha^{'}\lambda^{'}}{\lambda}-\frac{\alpha{\lambda^{'}}^{2}}{\lambda^{2}}+\frac{\alpha\lambda^{''}}{\lambda}+\alpha^{''}\left( \Psi\left( \sum_{j} y_{ij}+\alpha\right)-\Psi\left( \alpha\right) \right)+\alpha^{'}\left( \Psi^{'}\left( \sum_{j} y_{ij}+\alpha\right)-\Psi^{'}\left( \alpha\right) \right)-\frac{2\alpha^{'}\lambda^{'}}{\check{\mu}_{i}}-\alpha^{''}\log\check{\mu}_{i}-\lambda^{''}\check{\omega}_{i}+\frac{{\lambda^{'}}^{2}\check{\omega}_{i}}{\check{\mu}_{i}}+\left( \frac{\alpha^{'}-\lambda^{'}\check{\omega}_{i}}{\check{\mu}_{i}} \right)^{2}\sum_{j} \frac{\check{\upsilon}_{ij}{\mu_{ij}^{*}}^{2}}{\phi_{ij}^{*}}+\left( \frac{\alpha^{''}-\lambda^{''}\check{\omega}_{i}}{\check{\mu}_{i}}-\frac{2\alpha^{'}\lambda^{'}+2\lambda^{'}\check{\omega}_{i}}{{\check{\mu}_{i}}^{2}} \right)\sum_{j} \left( 1-\check{\upsilon}_{ij} \right)\mu_{ij}^{*}$$

$$\frac{\partial^{2}\tilde{l}_{i}\left( \boldsymbol{\beta,}\sigma^{2},\phi\right)}{\partial\phi\partial\sigma^{2}}=-\frac{\alpha^{'}-\lambda^{'}\check{\omega}_{i}}{\check{\mu}_{i}}\sum_{j} \frac{\left( 1-\check{\upsilon}_{ij} \right)\mu_{ij}^{*}}{\phi_{ij}^{*}}$$

$$\frac{\partial^{2}\tilde{l}_{i}\left( \boldsymbol{\beta,}\sigma^{2},\phi\right)}{\partial\phi\partial\beta_{s}}=-\check{\omega}_{i}\sum_{j} \frac{(1-\check{\upsilon}_{ij})\mu_{ij}^{*}x_{ijs}}{\phi_{ij}^{*}}-\frac{\check{\omega}_{i}\bar{x}_{is}}{\check{\mu}_{i}}\sum_{j} \frac{(1-\check{\upsilon}_{ij})\mu_{ij}^{*}}{\phi_{ij}^{*}}$$

$$\frac{\partial^{2}\tilde{l}_{i}\left( \boldsymbol{\beta,}\sigma^{2},\phi\right)}{\partial\sigma^{2}\partial\beta_{s}}\approx-\frac{\alpha^{'}\bar{x}_{is}}{\check{\mu}_{i}}+\frac{\lambda^{'}\check{\omega}_{i}\bar{x}_{is}}{\check{\mu}_{i}}+\left( \frac{\alpha^{'}}{\check{\mu}_{i}}-\lambda^{'}\check{\omega}_{i} \right)\check{\omega}_{i}\left( \sum_{j} \frac{\check{\upsilon}_{ij}{\mu_{ij}^{*}}^{2}x_{ijs}}{\phi_{ij}^{*}}+\frac{\bar{x}_{is}}{\check{\mu}_{i}}\sum_{j} \frac{\check{\upsilon}_{ij}{\mu_{ij}^{*}}^{3}}{\phi_{ij}^{*}} \right),$$

where $\Psi^{'}\left( x \right)=\frac{d^{2}\log(\boldsymbol{\Gamma}\left( x \right))}{dx^{2}}$ is the trigamma function, and $\frac{\partial^{2}\tilde{l}_{i}\left( \boldsymbol{\beta,}\sigma^{2},\phi\right)}{\partial\sigma^{2}\partial\beta_{s}}$ is approximated by plugging in $\sum_{j} \left( 1-\check{\upsilon}_{ij} \right)\approx0$.
